# Localized Rift Valley Fever Virus Persistence Depends on a High Transovarial Transmission Fraction

**DOI:** 10.1101/2023.10.27.564291

**Authors:** Melinda K. Rostal, Jamie Prentice, Noam Ross, Alan Kemp, Peter N. Thompson, Assaf Anyamba, Sarah Cleaveland, Claudia Cordel, Veerle Msimang, Petrus Jansen van Vuren, Daniel T. Haydon, William B. Karesh, Janusz T. Paweska, Louise Matthews

**Affiliations:** EcoHealth Alliance, New York, NY 10018 USA; School of Biodiversity, One Health and Veterinary Medicine, University of Glasgow, Glasgow G12 8QQ, UK; The Roslin Institute, University of Edinburgh, Edinburgh EH25 9RG, UK; Centre for Emerging Zoonotic and Parasitic Diseases, National Institute for Communicable Diseases, National Health Laboratory Service, Johannesburg 2192, South Africa; Epidemiology Section, Department of Production Animal Studies, Faculty of Veterinary Science, University of Pretoria, Onderstepoort 0110, South Africa; Universities Space Research Association, Columbia, MD 21046, USA; NASA Goddard Space Flight Center, Biospheric Sciences Laboratory, Greenbelt, MD 20771, USA; Geospatial Science and Human Security Division, National Security Sciences Directorate, Oak Ridge National Laboratory, Oak Ridge, TN 37830 USA; Department of Civil and Environmental Engineering, The University of Tennessee, Knoxville, Knoxville, Tennessee 37996 USA; ExecuVet PTY LTD., Bloemfontein 9301, Free State, South Africa; Australian Centre for Disease Preparedness, CSIRO-Health and Biosecurity, Geelong, Victoria, 3220, Australia

**Keywords:** Rift Valley fever, RVF, transovarial transmission, mechanistic model, vaccination

## Abstract

Rift Valley fever virus (RVFV) has spread beyond continental Africa and threatens to follow West Nile, chikungunya and Zika viruses into the Americas. Its impact in new localities and the capacity to control future outbreaks, depends on whether and how RVFV persists at small spatial scales. Transovarial transmission (TOT) is hypothesized as an important mechanism for local persistence, yet its role in RVFV ecology remains poorly understood. We examine whether RVFV can persist locally via TOT while maintaining a realistic seroprevalence pattern of interepidemic and epidemic transmission. We developed a mechanistic, compartmental model of RVFV dynamics within a single host (sheep) and two vector (mosquito) populations, driven by temperate climatic factors. Decades-long persistence was possible in our simulations, which generally captured the observed outbreak patterns in central South Africa with a mean annual seroprevalence (∼23%) within the range reported during interepidemic periods (5-40%). Persistence was only possible with a substantial TOT fraction and over a narrow range of parameters. The basic reproduction number (*R*_0_) was close to one at mean vector population sizes, suggesting a relatively limited expansion of the infected vector population during outbreaks. This limited expansion provides the system with the flexibility to support both low-level transmission and large outbreaks and, counterintuitively, large outbreaks resulted in smaller infected *Aedes* egg populations. This has important consequences for control: low-level vaccination may prevent large outbreaks without eliminating RVFV and local control efforts may be most effective immediately following an outbreak, suggesting elimination may be possible after emergence in temperate regions.

## Introduction

Arboviruses have emerged and re-emerged globally during the past 50 years (1), including chikungunya, West Nile and Zika viruses (1, 2). A better understanding of viral ecology (e.g. over-wintering mechanisms (3)) could have allowed faster detection and response during the initial epidemics, reducing the magnitude of the outbreaks and the risk of the viruses establishing endemicity. Rift Valley fever virus (RVFV) has spread across Africa and emerged in new regions outside of Africa. Important gaps in understanding RVFV persistence and maintenance mechanisms limit our ability to predict and target mitigation efforts against outbreaks and expansion.

First reported in Kenya in 1931 (4), RVFV was detected in South Africa (1950; (5)) and subsequent outbreaks indicate endemicity throughout Africa (6). Specifically, South Africa reported widespread outbreaks in 1950-1951, 1974-1975 and 2010-2011 and multiple, isolated outbreaks during intervening years (7). In 1990, RVFV emerged in Madagascar, the first report outside continental Africa (8), with further emergence in Saudi Arabia/Yemen (2000; (9)) and the Comoros (2007 (10)). This pattern of emergence suggests there is a high risk of global RVFV spread. However, the underlying mechanisms of RVFV persistence have not been identified, despite a significant body of work focused on RVFV ecology.

Widespread and severe outbreaks of Rift Valley fever (RVF) manifest as abortion storms and high neonatal mortality in ruminant livestock resulting in significant economic losses (11). Clinical disease in people ranges from mild fever and myalgia to rare hemorrhagic fever cases (12). Outbreak periods are interspersed with interepidemic periods lasting 2-25 years (13). The mean seroprevalence estimate from interepidemic ruminant studies conducted across Africa was 15% (range: 0-44%) (13). During interepidemic periods, there is evidence of RVFV circulation in livestock at a sufficiently low frequency that cases often remain undetected (14, 15). Wildlife can also be infected, and seroconversions during interepidemic periods suggest that some species may also support low-level RVFV interepidemic circulation, though the role they play in maintenance remains unknown (16).

After RVFV was detected in larval and unfed *Aedes mcintoshi*, these floodwater mosquitoes were proposed to be a primary reservoir of RVFV and to maintain it via transovarial transmission (TOT), during dry periods (17). More recently, RVFV antigen was detected in the eggs of *Ae. mcintoshi* collected in Kenya (18). Due to difficulties in establishing floodwater *Aedes* breeding colonies, laboratory estimation of RVFV TOT fraction, the proportion of infected eggs laid by an infected female mosquito, has yet to be measured in African mosquitoes (19). However, a recent study provided the first experimental evidence of RVFV TOT, identifying transovarial transmission in 2-10% of progeny of infected females of the North American mosquito, *Culex tarsalis* (20). To date, there is no field evidence that *Culex* spp. can pass RVFV transovarially, though they can transmit RVFV horizontally. Possible overwintering mechanisms for RVFV include TOT, animal movement (metapopulation dynamics), or the survival of infected adult mosquitoes, as has been reported for other pathogens (3). Transovarial transmission by floodwater *Aedes* is likely an important mechanism of RVFV overwintering in endemic regions that face dry conditions (21), supporting low-level, annual transmission to livestock and wildlife interspersed with large outbreaks (22). Yet, no mechanistic model has demonstrated the feasibility of this persistence mechanism at a localized scale (e.g., a small, seasonal wetland or pan), while producing realistic host infection dynamics.

We examine the mechanisms of decadal, localized RVFV persistence that reflects historical climatic and epidemic conditions of a regional epicenter for widespread RVF outbreaks, the Free State Province, South Africa. We developed a SEIR model of sheep (SIR), *Culex* (SEI, horizontal transmission) and floodwater *Aedes* (SEI, TOT and with a univoltine life cycle - the eggs hatch during one period per year). Specifically, we examine whether interepidemic persistence is possible at this local scale, examine the role of TOT in persistence, identify other key parameters in the maintenance of RVFV, estimate the reproduction number (*R*_0_) and examine vaccination implications.

## Results

### Decadal, localized persistence is possible

Our simulations indicated that RVFV can persist for over thirty years within a localized area or pan producing realistic outbreak and infection patterns (Figure 1) over a narrow range of parameters (Figure 2, orange and black striped region). The exemplar simulation generated small annual RVF outbreaks with three large outbreaks (infecting >25% of the flock) occurring in 1987, 1999 and 2009 (Figure 1A) that were associated with a higher ratio of infected *Culex* to infected *Aedes* (Figure 1B, S3). Two of these large outbreaks aligned well to reported outbreaks in South Africa during the timeframe, the largest of which was in 2010-2011 (Figure 1A). The population of infected *Aedes* eggs remained stable throughout the simulation (Figure 1C) and the mean proportion of infected *Aedes* eggs was 0.003 (Table S1). The exemplar simulation had a low mean annual proportion of infected adult vectors, ranging from 0-0.04 (Table S1), and the mean annual seroprevalence was 22.9% (median: 16.3%; range 10.5-58.6%; Table S1). Excluding the three large outbreaks, the average incidence was 7 per 100 hosts per year. The calculated *R*_*0*_ was 1.05 at the mean population sizes of *Aedes* and *Culex*, increasing up to 1.9 at the mean of the annual peak population sizes for the exemplar simulation. *R*_*0*_ was most sensitive to the *Aedes* and *Culex* bite and mortality rates (Figures S4 & S5).

**Figure 1.**
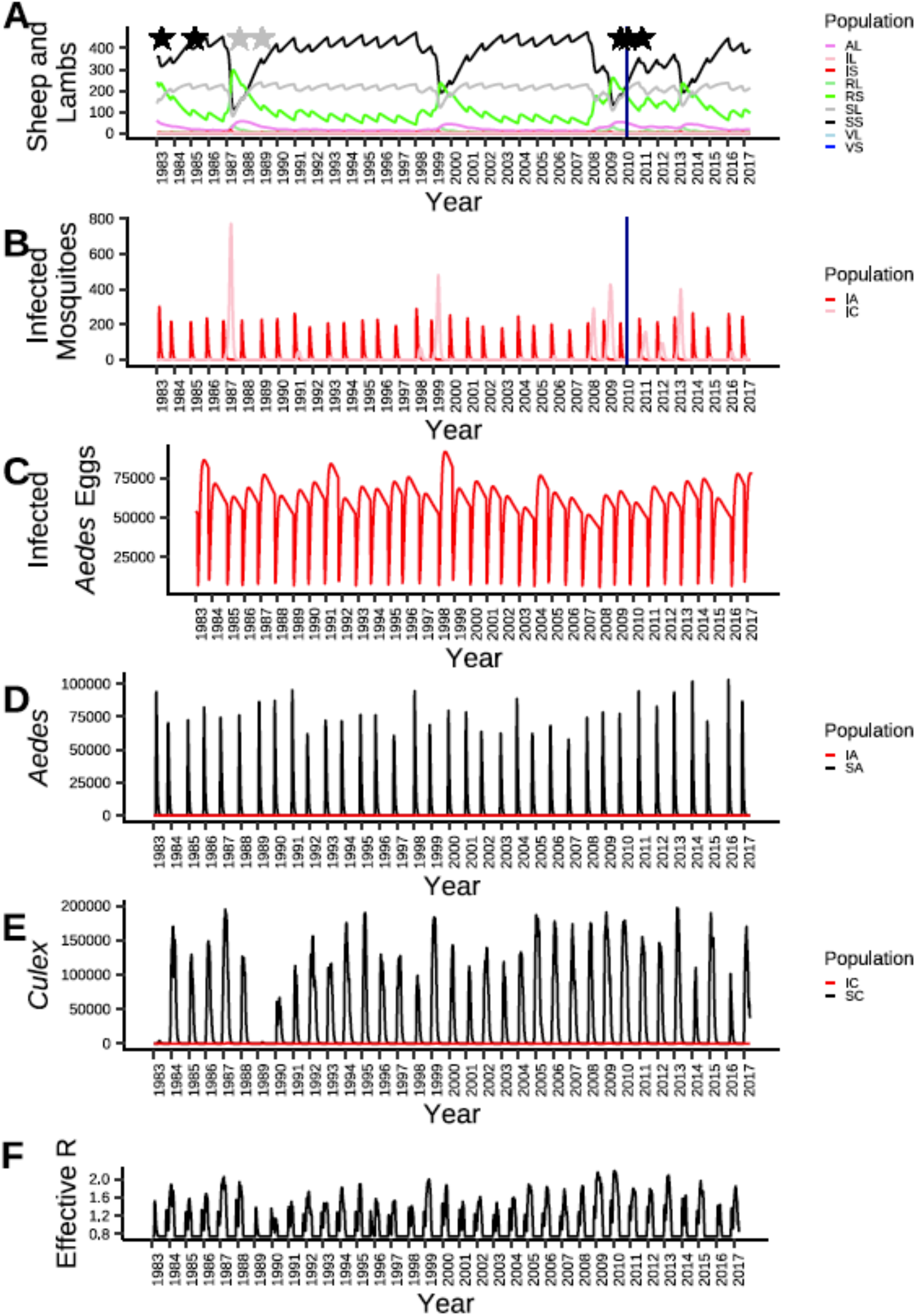
The full simulation of RVFV infection dynamics with two vectors (Aedes and Culex) in a single host system run for 34 years. Annual circulation is evident with several small outbreaks over the years and three very large outbreaks (one of which occurred in 2009). A) The simulated populations of susceptible (black/grey), infected (red/pink) and recovered (green/light green) sheep/lambs. The dark blue line indicates the approximate time of the 2010 outbreak. The black stars indicate year where there was a recorded RVF outbreak in the Central Plateau portions of the Free State and/or Northern Cape. The gray stars indicate a year in which an RVF outbreak was recorded but it is unknown where in South Africa the outbreak occurred (7). B) The populations of infected Aedes (red) and Culex (pink) mosquitoes are shown and the 2010 outbreak is marked by the dark blue line. C) The number of infected Aedes eggs in the environment is indicated in red over time. D) The total populations of susceptible (black) and infected (red) Aedes mosquitoes are shown over time. E) The total populations of susceptible (black) and infected (red) Culex mosquitoes are shown over time. F) Changes in the effective reproduction number as the simulation proceeds. AL = lambs with maternal antibodies; IL, IS, IA, and IC = infected lambs, sheep, Aedes, and Culex, respectively; RL and RS = recovered lambs and sheep, respectively; SL, SS, SA, and SC = susceptible lambs, sheep, adult Aedes and adult Culex, respectively. VL and VS = vaccinated lambs and sheep, respectively; however, the plotted simulation did not include any vaccinated animals.

**Figure 2.**
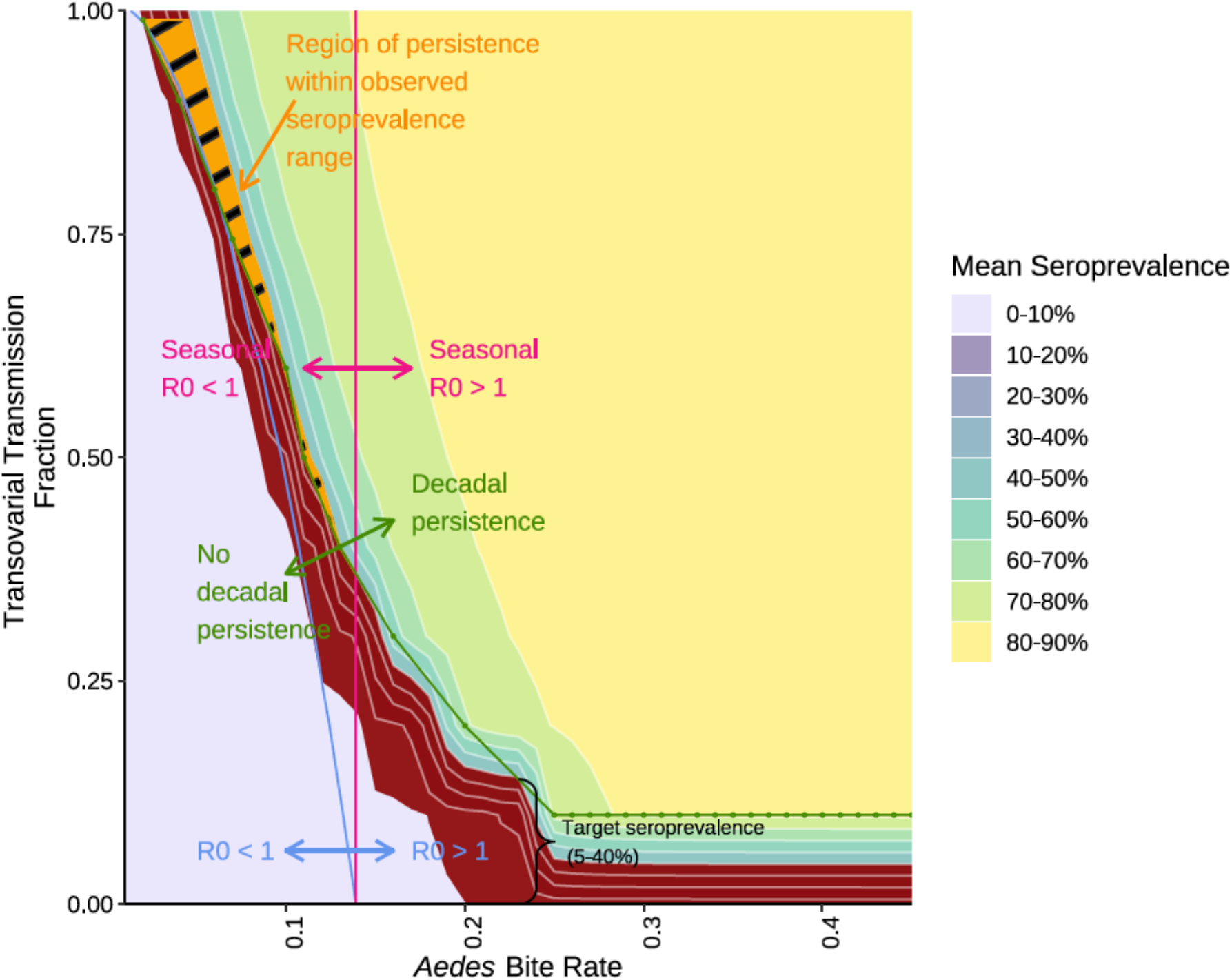
Contours of mean seroprevalence as TOT fraction and Aedes bite rate are varied. The target seroprevalence range (5-40%) is highlighted in red. For each TOT fraction simulated (intervals of 0.1), the lowest Aedes bite rate that supported decadal (34-year) persistence was plotted (green line), such that parameter value combinations in the space above the green line support decadal persistence. The contour line representing interannual ***R***_***0***_ = 1 is shown in blue. The system is capable of short-term RVFV persistence (***R***_***0***_ *≥* 1) in the parameter space at or above/to the right of the blue line. To evaluate the contribution of TOT to ***R***_***0***_ we identified the parameter space where the seasonal ***R***_***0***_ was equal to 1 (calculated with zero contribution of TOT; pink line) for each simulation, the space to the right of the pink line is where the seasonal ***R***_***0***_ is greater than one. We see that with TOT, decadal persistence is possible, though it results in seroprevalences that are higher than is observed in nature. The orange and black region represents the parameter space that supported decadal persistence with ***R***_***0***_ greater than one within the targeted seroprevalence. This only occurs when the seasonal ***R***_***0***_ is less than one.

### Transovarial transmission, a key parameter for RVFV persistence

In the absence of TOT, the virus became locally extinct after one year of simulation (Figure S6A & B). In a system with no horizontal transmission (i.e., no viral expansion in the host or *Culex* populations), RVFV became extinct in the host population after seven years, but infected *Aedes* adults were present for up to a decade (Figure S6C & D). Viral persistence increased gradually as TOT fraction increased up to 0.6 (commensurate with up to nine years of persistence) beyond which the system rapidly changed to support persistence for the entire 34-year period (Figure S7B). In our model, TOT fractions between 0.67 and 0.81 supported mean seroprevalence levels of 5-41% (Figure S7A).

In our sensitivity analysis, all six outcome factors (RVFV persistence, mean annual seroprevalence, mean outbreak size, mean outbreak length and maximum outbreak size) were sensitive to TOT fraction (Figure 3). Viral persistence and the mean seroprevalence in the 34-year system were very sensitive to small changes in the *Aedes* transmission parameters, bite rate and the external incubation rate (Figure S7). As we varied key parameter(s), we saw rapid switching of behaviors from systems with low seroprevalence and short-term persistence to systems with high seroprevalence and long-term persistence (Figures S7 and S8). Among the 4000 runs in the sensitivity analysis, 53% resulted in RVFV extinction and 37% resulted in unrealistically high seroprevalences.

**Figure 3.**
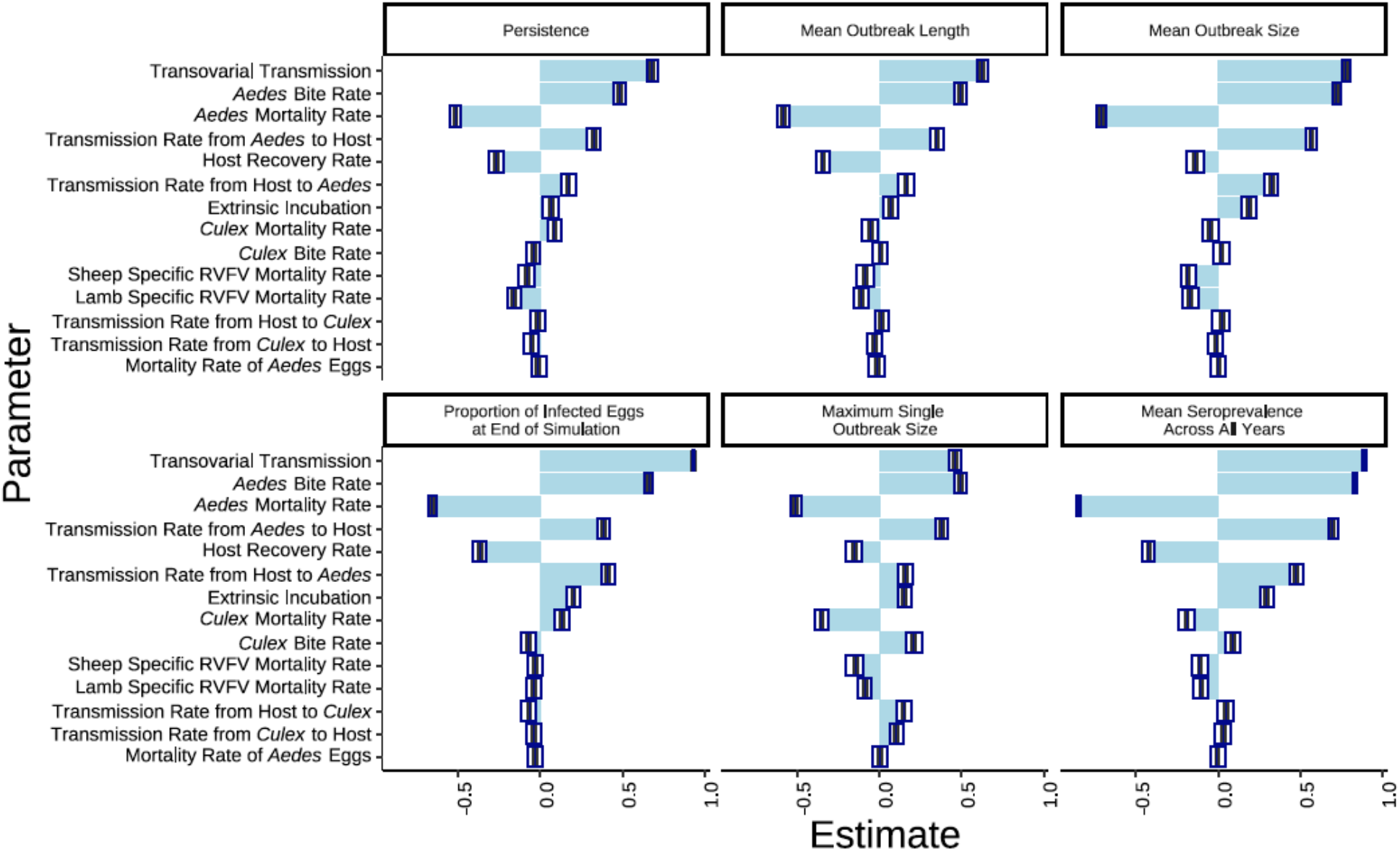
Tornado plot of the partial coefficient correlation estimates and 99% confidence interval indicating the sensitivity of each outcome factor of interest to the selected parameters in the model. The box plot shows the 99% confidence interval, and the blue bar shows the coefficient estimate. The larger the absolute value of the estimate was, the more sensitive that outcome factor is to a specific parameter.

Decadal persistence at target (5-40%) seroprevalence ranges (Figure 2, orange hatched region) required *R*_*0*_ > 1 and seasonal *R*_*0*_ < 1 (where seasonal *R*_*0*_ is calculated as *R*_*0*_ without TOT to capture the within year dynamics; Figures 2 and S8). When requiring target seroprevalence and RVFV persistence, *R*_*0*_ was only consistently above unity at a high TOT fraction (>0.6), though persistence was possible for occasional simulations with a TOT of 0.4-0.5 (Figure 2). Seasonal *R*_*0*_ < 1 indicates relatively little expansion of the infected vector population during outbreaks. Simulations where the seasonal *R*_*0*_ was above unity and within the target seroprevalence resulted in multiple large outbreaks (*R*_*0*_>1 during outbreak years), which drove RVFV extinct.

### Seasonal R_0_ and vector dynamics drive outbreaks

In our simulations, small numbers of susceptible *Culex* survived most winters, though no infected *Culex* survived a season. Large outbreaks (seasonal *R*_*0*_ > 1) occurred when there was RVFV expansion within the *Culex* population (Figures 1B, S3 and S9). These dynamics are exemplified by simulations with a 25% lower *Culex* mortality rate that resulted in RVFV extinction (Figure S9, left). While the system was resilient to a single large outbreak resulting in high host immunity (Figure 4D), multiple large outbreaks (resultant of RVFV expansion in the *Culex* vector), eventually led to RVFV extinction in the host population (Figure S9A, left) and a reduction in the number of infected *Aedes* eggs over time (Figure S9C, left). Similarly, the sensitivity analysis outcome that was most affected by the *Culux* parameters was the maximum single outbreak size (Figure 3).

**Figure 4.**
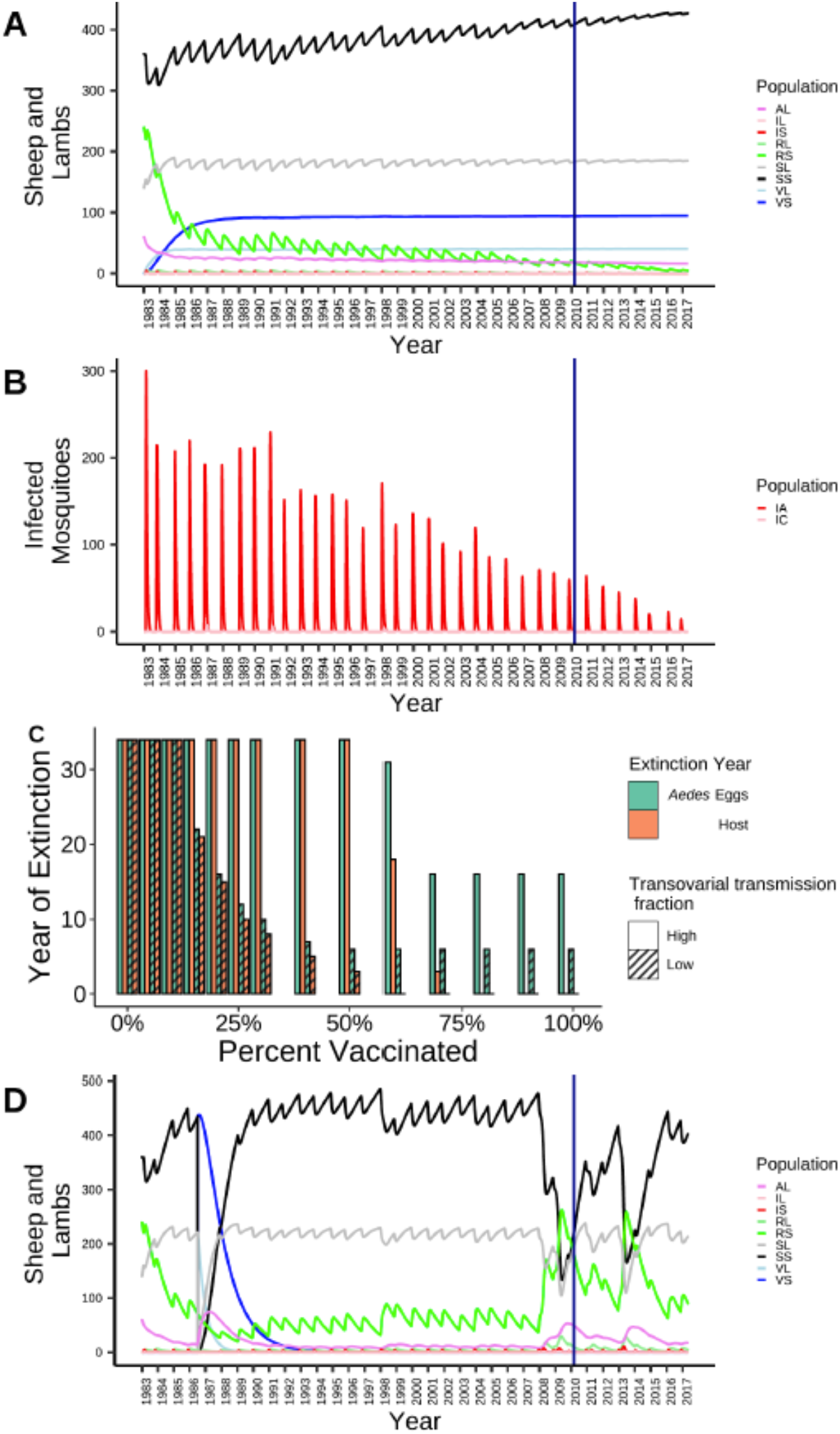
Simulation of the RVFV dynamics when 18% of susceptible lambs are vaccinated for A) the sheep and lamb populations and B) the infected Aedes egg population. C) The level of vaccination the flock is maintained at versus the number of years until extinction in the host and *Aedes* egg populations at low (0.75) and high (0.9) transovarial transmission fractions. D) In a simulation where 99% of the flock is vaccinated in 1986, RVFV persists in the infected Aedes eggs and infections continue until the large outbreak in 2009 and beyond.

### Implications for vaccination as a mitigation tool

Under vaccination of the host population, infection in the vector population was maintained after extinction in the host population, with a greater difference in extinction times for higher TOT (Figure 4C, Figure S10). To drive RVFV to extinction required vaccination levels of at least 18% to be maintained in the host for 27 years (Figure 4A). At vaccination levels of 50% and TOT of 0.75 extinction could be achieved in 3 years when TOT is 0.75, but sufficient infection remained in the vector population to initiate outbreaks for 6 years. At higher TOT (0.9), infection would persist in host and vector (Figure 4C).

Burst vaccination, vaccinating 99% of susceptible sheep and lambs during a 7-day period during winter, also required several years to overcome TOT. Simulating burst vaccination immediately after a large outbreak or earlier in the simulation when the infected Aedes egg population was less established drove RVFV to extinction faster (Figure S11).

## Discussion

As RVFV spreads beyond Africa, it is critical to understand mechanisms of transmission to prevent RVFV establishment in new regions. We simulated local persistence of RVFV, which reproduced historical outbreak patterns and resulted in a realistic seroprevalence pattern Decadal persistence in this closed system under temperate environmental conditions was dependent on TOT and limited expansion in the vector populations, though it was very sensitive to multiple key parameters. Multiple large outbreaks resulting in rapid expansion of RVFV in the host population and driven by *Culex* led to a high seasonal *R*_*0*_ and reduced persistence by driving RVFV extinct within the *Aedes* egg population. To prevent long-term persistence, a consistent and annual vaccination regimen, or at the very least several years of vaccinating 99% of the host population, is necessary. Vaccination may be most effective in reducing the population of infected *Aedes* eggs if used immediately following a large outbreak.

Immediate response and largescale vaccination after RVFV emergence may be able to prevent establishment if RVFV emerges in a new temperate region.

### Long-term, localized persistence is possible

Our simulations suggest that RVFV can persist locally for decades. While not predictive, our model produced a realistic pattern of outbreaks with two of the three large outbreaks occurring close to times when outbreaks were reported by the Free State or an unspecified location in South Africa (7). The simulated outbreak in 1987 was larger than expected from the literature (7), this may be result of initial conditions (in 1983). The large, simulated outbreak in 2009 fits reasonably well to the timing of the large 2010-2011 outbreaks in the Free State. This suggests that the model well-approximates the mechanism for localized RVFV persistence.

The mean annual seroprevalence simulated was 22.9% (range 10.5-58.6%), which is within the range of reported ruminant seroprevalence estimates across Africa (13). The range overlaps with the confidence interval of a recent seroprevalence estimate in unvaccinated ruminants in the Free State (23). It is important to evaluate whether the model can reproduce the observed pattern of low interepidemic transmission interspersed with large outbreaks. Most other simulation models either simulated a high proportion of immune/recovered animals over time (24, 25) or only presented the infectious host dynamics to demonstrate RVFV infection periodicity without presenting the recovered or susceptible hosts dynamics (26, 27). The few studies that do simulate interepidemic periods with low seroprevalence, are at a regional scale (28), path B), (29, 30) and/or incorporate movement using a metapopulation structure (29, 31). A metapopulation system may be able to support RVFV maintenance as ruminant movement along livestock trading routes has been implicated in RVFV spread (32-34) and maintenance. However, isolated RVF outbreaks have been observed (7) and likely contribute to observed spatially patchy patterns of seroprevalence (35), which suggests local persistence occurs. If the TOT fraction is lower than the range we found was required for local persistence, a metapopulation model or other reservoir mechanism would be required.

### Transovarial transmission, a key parameter for RVFV persistence

Our analysis suggests that TOT is likely a key contributor to long-term RVFV persistence in a system where temperature and/or rainfall patterns make the survival of overwintering, RVFV-infected mosquitoes unlikely. Transovarial transmission is an important contributor toward *R*_*0*_ surpassing unity (discussed below) and provides a mechanism for overwintering. Other models have also suggested that TOT is critical in ecosystems with a prolonged dry period (21, 24). However, simulation of RVFV dynamics in more favorable climatic conditions may not require TOT for RVFV persistence (21, 28, 29)). Iacono et al.’s (28) path A model suggested that TOT of RVFV was not required, although this simulation resulted in extremely high interepidemic seroprevalence (approximately 80-90%). On the regional scale (based on the water surface area available in Kenya), they were able to simulate large RVFV outbreaks interspersed with interepidemic periods with a low seroprevalence (path B) when environmental conditions (mean annual temperatures 18-22°C and mean water surface area 3,000-4,000 m^2^) resulted in “unstable persistence” of RVFV, included environmental transitions between systems that did and did not require an overwintering mechanism and resembled chaotic behavior. Regions in both Kenya (e.g., Ijara) and South Africa have prolonged dry seasons, which in South Africa is coupled with a temperate ecosystem. Our simulation suggests that TOT is likely to be critical to support RVFV persistence under these conditions.

Our model was highly sensitive to multiple parameters, including *Aedes* and *Culex* bite rates and transmission parameters. Small changes to these parameters led either to extinction or to a persistent, unrealistically high, host seroprevalence. Our model’s sensitivity suggests that while RVFV may persist in some pans, RVFV extinction is possible in other pans depending on how these traits/rates vary with host and vector population sizes and composition. The proportion of sensitivity analysis simulations in which RVFV went extinct supports conditional and heterogenous persistence. Future investigation with a stochastic model could estimate the local extinction rate, though it would likely result in a smaller parameter space in which observed dynamics would arise. Favier *et al*. (36) suggested that, although RVFV may become extinct at local scales, it could be maintained within a network of local pans by the movement of livestock between seasonally inundated pans. Sumaye et al. (31) also used movement to simulate RVFV dynamics with a low interepidemic seroprevalence. Future extensions of our current model to the metapopulation-level could examine persistence more thoroughly.

A high TOT fraction (generally >0.6) was required to support persistence in our system. Sumaye *et al*. (31) also used a high TOT fraction (0.5) to simulate RVFV persistence with a comparable seroprevalence. These results challenge the common, unsupported, perception that the RVFV TOT fraction is low (19). This perception may have developed because of the relatively low detection rate of RVFV in adult mosquito populations (e.g. 0.001 reported on farms following RVF outbreaks in 1974-5 (37)). However, our simulations suggest that even with a high TOT fraction, the mean proportion of infected adults remains relatively low (0-0.04). A high TOT fraction for mosquito viruses is biologically plausible, as *Culex* flavivirus is solely maintained by a TOT fraction of 0.97 (38). The only study to experimentally demonstrate RVFV TOT estimated it to be 0.02-0.10 in a species of *Culex* that has never been exposed to RVFV in nature (20). A natural vector of RVFV would likely have evolved a higher TOT fraction. Future establishment of floodwater *Aedes* spp. breeding colonies to assess TOT fraction is needed.

Our sensitivity analysis indicated that TOT had a large effect on all outcome factors. Though *R*_*0*_ had little sensitivity to changes in TOT fraction, we found that TOT makes an important contribution to *R*_*0*_ reaching unity in simulations with long-term persistence and are within the target seroprevalence range. This finding holds even when varying parameters that *R*_*0*_ is very sensitive to, e.g., *Aedes* bite rate. We saw consistent long-term persistence within the target seroprevalence range when TOT was above 0.6. In these scenarios, the seasonal *R*_*0*_ was less than one, indicating the important contribution of TOT. When the target seroprevalence was achieved and the seasonal *R*_*0*_ (calculated with zero TOT fraction) exceeded one there were large RVF outbreaks without long-term persistence. Thus, achieving long-term persistence and observed seroprevalences appears to require limited seasonal RVFV spread and an overwintering mechanism via a high TOT to boost *R*_*0*_ above unity. Previous analyses have not simultaneously examined *R*_*0*_ with multi-year persistence, which may explain why many have not found *R*_*0*_ to be sensitive to TOT (24, 26). Given that American *Culex* spp. are capable of TOT, if American *Aedes* spp. have a higher TOT fraction, TOT could be an overwintering mechanism to sustain RVFV maintenance upon introduction to the Americas (20).

### Seasonal R_0_ and vector dynamics drive outbreaks

The interactions between the *Aedes* and *Culex* mosquitoes were important drivers of RVFV outbreak dynamics. Though our sensitivity analysis indicated that decadal persistence and the infection dynamics were primarily driven by *Aedes* traits, with little sensitivity to *Culex* traits, we found that *Culex* bite and mortality rates were important drivers of outbreak size (contributing to seasonal *R*_*0*_). *R*_*0*_ was similarly sensitive to *Culex* traits, especially high *Culex* bite rates, in other simulation studies (24, 39-41).

Low-level interepidemic transmission of RVFV requires that there be little expansion of RVFV in the vector and host populations. The large RVF outbreaks were driven by *Culex* and resulted in rapid expansion within the host population (seasonal *R*_*0*_> 1), while still maintaining a limited expansion within the much larger population of adult *Aedes* and *Culex* mosquitoes (maximum proportion of infected vectors was ∼4% within each population), which is in line with the low detection rate reported in mosquitoes during RVF outbreaks (37). Larger outbreaks induced high rates of host immunity, which counterintuitively led to declines in the infected *Aedes* egg population. The population size of infected *Aedes* eggs at the start of the season is likely important for a system where persistence is unstable.

Trait variation within the community of vectors and whether overwintering is required may explain the variation in interepidemic period duration among regions (13). Cavalerie *et al*. (21) estimated the horizontal vector transmission fraction based on the observed relative abundance of different mosquito species in Mayotte and their laboratory-determined transmission fractions. Thus, the potential for widely variable *Culex* population sizes and transmission, bite and demographic rates based on local vector communities may determine whether RVFV can persist via a given TOT fraction. Our model did not support continued growth of the *Culex* populations during most winters, though small populations of susceptible *adult Culex* survived. These overwintering *Culex* did not support RVFV.

### Implications for vaccination as a mitigation tool

The effect of TOT on RVFV persistence has implications for vaccination strategies. A low level of vaccination (18%) could drive RVFV to extinction; however, a high TOT fraction requires maintenance of this vaccination level for decades before extinction in this closed system. Higher annual vaccination levels drove RVFV extinct more rapidly. Given the importance of TOT, persistence of RVFV is only partially dependent on expansion in the host population and it can be maintained solely by the *Aedes* for several years before being driven to extinction. Higher levels of vaccination would be necessary if wildlife reservoirs or metapopulation dynamics contribute to RVFV maintenance. Even within a highly vaccinated (99%) host population, our simulated infected egg population sufficiently persisted for approximately six years and would be capable of initiating an outbreak if all immune hosts were removed.

To reduce or eliminate the local population of infected *Aedes* eggs, high vaccination levels may have a larger impact when implemented immediately following a large outbreak that stimulated high levels of host immunity. These strategies have the potential to reduce or eliminate the population of infected *Aedes* eggs within a local pan used exclusively by livestock. Though not evaluated here, pesticides use in a pan may additionally reduce the population of infected *Aedes* eggs, especially when used in conjunction with vaccination after an outbreak or any cause of reduced *Aedes* egg populations (e.g., drought). This would reduce or eliminate undiagnosed, interepidemic infections and prevent endemic RVF outbreaks at that pan, even in the presence of abnormally high precipitation and flooding. Applying this to RVFV emergence, it suggests that if intensive control measures (e.g., 99% vaccination of nearby livestock) are maintained for several years after an incursion of RVFV, it may be possible to eliminate RVFV before it is fully established.

## Materials and Methods

We developed a deterministic, ordinary differential equation (ODE)-based compartmental model with a single livestock host (sheep) and two mosquito vectors, forced by climatic parameters (Figure S1; Table S2; see Supplemental Information for full methods and model description). Specifically, we used a susceptible (S), infected (I) and recovered (R) system (SIR) to simulate the numbers of sheep and lambs (including compartments for passive immunity and vaccination) and a susceptible, exposed (E) and infected system (SEI) for the *Aedes* and *Culex* mosquitoes (Eqs. S1-S30). Our system included TOT in *Aedes* and amplification through horizontal transmission by *Culex* which continue to hatch throughout the season, maintaining the natural succession between these vectors (17, 42, 43). To simulate the typical interepidemic RVFV conditions associated with successive below-average rainfall seasons, we assumed a univoltine life cycle in the transovarially transmitting vector. We used 34 years (1983-2017) of rainfall data from Bultfontein (latitude: −28.40, longitude: 26.26) extracted from African Rainfall Climatology (ARC) data records (44, 45) and temperature data from the nearest South African Weather Service stations (latitude: −28.95, longitude: 26.33) in Free State, South Africa (Figure S2). The climate data were used to drive vector hatching and development and mortality rates ((46, 47); Tables S3 and S4; Eqs. S31-S33). The parameter values (Table S5) were taken from the literature, where available, and via expert opinion except for the values of four parameter groups (vector carrying capacities and bite rates, TOT fraction and *Aedes* egg mortality rate). We used a Latin hypercube to vary these parameters in 80,000 simulations. Each simulation was assessed using six criteria (Supplemental Information). The simulation that met the criteria and most closely resembled the historical pattern of RVF in the Free State was selected. The parameter values from this simulation were used for the exemplar and other simulations (Table S5). We used a conservative measure of persistence, sustaining at least one infection in the host population per season (September-August) for each of the 34 years of the simulation.

We conducted a sensitivity analysis of selected parameters (Table S6) using 4000 draws from a Latin hypercube (48, 49) (Supplemental Information). Each parameter was evaluated based on its effect on six outcomes: persistence, mean annual seroprevalence, the proportion of infected *Aedes* eggs at the end of the simulation, mean outbreak length, mean outbreak size and maximum outbreak size. Annual estimates were summarized over the mosquito growth season in South Africa (September-August). The results were analyzed using a partial coefficient correlation (PCC) analysis (48).

*R*_*0*_ was numerically estimated at the mean and peak vector and host population sizes using a next generation matrix (NGM) approach (50) (Eq. S34-36). The effective reproduction number (*R*_”_) was estimated using the susceptible host and vector populations at each timestep. We also conducted a sensitivity analysis of ***R***_0_ to varying parameters and investigated long-term persistence as it related to ***R***_0_ and the seasonal ***R***_0_ (calculated by excluding the contribution of TOT to ***R***_0_). See Supplemental Information.

## Acknowledgments

We want to acknowledge the data provided by the South African Weather Service, which provided the temperature data that was used to drive the model. The project depicted is sponsored by the U.S. Department of Defense, Defense Threat Reduction Agency. The content of the information does not necessarily reflect the position or the policy of the federal government, and no official endorsement should be inferred.

