## Supplemental methods, figures and tables for "Localized Rift Valley Fever Virus Persistence Depends on a High Transovarial Transmission Fraction"

**Supporting Information Text**

**Detailed methods and materials**

We developed a deterministic, ordinary differential equation (ODE)-based model with two mosquito vectors and a single host driven by realistic climate forcing. Specifically, to drive the vector population dynamics and simulate the RVFV dynamics during a 34-year period, we used rainfall and temperature data (described below) from a representative farm in central South Africa. The vector-borne component is conceptually based on the MacDonald malaria model (1) with the addition of transovarial transmission in one vector species, more development stages and climate forcing to drive the population dynamics of both vector species. A single, large sheep flock situated within a single pan in central South Africa was selected as the appropriate epidemiological unit. The transovarial *Aedes* vector was modelled with a univoltine life cycle and with the *Culex* vector hatching shortly after the *Aedes*. We used a susceptible (S), infected (I) and recovered (R) model in one host species with two life stages: adult sheep (S) and lambs (L) and a susceptible (S), exposed (E), infected (I) model for the two vector species: *Aedes* mosquitoes (A) and *Culex* mosquitoes (C) with multiple life stages (see Table S1 for all state variable definitions; Figure S1). *Aedes* can transmit RVFV both transovarially and horizontally, while *Culex* can only transmit RVFV horizontally. Summaries of the number of hosts or vectors infected were calculated over a yearly timeframe that represents the hatching season in South Africa (September 1 – August 31).

**Model Equations**

The parameters for the model are given in Table S4. The equations describing the sheep (which were compartmentalized as susceptible, SS, infected, IS, recovered, RS, or vaccinated, VS) and lamb populations (which were compartmentalized as susceptible, SL, infected, IL, recovered, RL, those with maternal immunity, AL, or vaccinated, VL) are as follows:

$$\frac{dSS}{dt} = g \cdot SL - \mathrm{sold}_S \cdot SS - \theta_{SLC} \cdot SS \cdot IC - \theta_{SLA} \cdot SS \cdot IA - \mu_S \cdot SS \quad \text{(S1)}$$

$$\frac{dIS}{dt} = g \cdot IL - \mathrm{sold}_S \cdot IS + \theta_{SLC} \cdot SS \cdot IC + \theta_{SLA} \cdot SS \cdot IA - \mu_S \cdot IS - \sigma \cdot IS - \rho_S \cdot IS \quad \text{(S2)}$$

$$\frac{dRS}{dt} = g \cdot RL - \mathrm{sold}_S \cdot RS - \mu_S \cdot RS + \sigma \cdot IS \quad \text{(S3)}$$

$$\frac{dVS}{dt} = g \cdot VL + vax \cdot SS \cdot vax_{scheme.s} - \mathrm{sold}_S \cdot VS - \mu_S \cdot VS \quad \text{(S4)}$$

$$\frac{dSL}{dt} = b_L \cdot \left(1 - \frac{NS}{NL_{max}}\right) \cdot SS + \omega_{MA} \cdot AL + \mathrm{buy}_L - g \cdot SL - \theta_{SLC} \cdot SL \cdot IC - \theta_{SLA} \cdot SL \cdot IA - \mu_L \cdot SL - \mathrm{vax} \cdot SL \quad \text{(S5)}$$

$$\frac{dIL}{dt} = \theta_{SLC} \cdot SL \cdot IC + \theta_{SLA} \cdot SL \cdot IA - g \cdot IL - \mu_L \cdot IL - \sigma \cdot IL - \rho_L \cdot IL \quad \text{(S6)}$$

$$\frac{dRL}{dt} = \sigma \cdot IL - g \cdot RL - \mu_L \cdot RL \quad \text{(S7)}$$

$$\frac{dAL}{dt} = b_L \cdot \left(1 - \frac{NS}{NL_{max}}\right) \cdot (RS + ((1 - Abort) \cdot IS + VS)) - \omega_{MA} \cdot AL - \mu_L \cdot AL \quad \text{(S8)}$$

$$\frac{dVL}{dt} = vax \cdot SL \cdot vax_{scheme.l} - g \cdot VL - \mu_L \cdot VL \quad \text{(S9)}$$

The equations for the *Aedes* free-living stages are given, followed by the equations for the egg stages. These were compartmentalized as susceptible and infected larvae and pupae, SALP and IALP, young adults (prior to first feeding), SAY and IAY, adults, SA and IA, eggs that are mature enough to hatch, SAE and IAE, and eggs that were laid this season and are not mature enough to hatch yet, $SAE_{new}$ and $IAE_{new}$, respectively. There is also a compartment for exposed adults, EA.

$$\frac{dSALP}{dt} = bh_A \cdot \mathrm{End}_A \cdot SAE \cdot \left(1 - \frac{NALP}{NALP_{max}}\right) - \mathrm{dev}_{ALP} \cdot SALP - \mu_{ALP} \cdot SALP \quad \text{(S10)}$$

$$\frac{dSAY}{dt} = \mathrm{dev}_{ALP} \cdot SALP - \mathrm{wait}_A \cdot SAY - \mu_A \cdot SAY \quad \text{(S11)}$$

$$\frac{dSA}{dt} = \mathrm{wait}_A \cdot SAY - \theta_{ASL} \cdot SA \cdot (IL + IS) - \mu_A \cdot SA \quad \text{(S12)}$$

$$\frac{dEA}{dt} = \theta_{ASL} \cdot SA \cdot (IL + IS) - \varepsilon \cdot EA - \mu_A \cdot EA \quad \text{(S13)}$$

$$\frac{dIALP}{dt} = bh_A \cdot \mathrm{End}_A \cdot IAE \cdot \left(1 - \frac{NALP}{NAE_{max}}\right) - \mathrm{dev}_{ALP} \cdot IALP - \mu_{ALP} \cdot IALP \quad \text{(S14)}$$

$$\frac{dIAY}{dt} = \mathrm{dev}_{ALP} \cdot IALP - \mathrm{wait}_A \cdot IAY - \mu_A \cdot IAY \quad \text{(S15)}$$

$$\frac{dIA}{dt} = wait_A \cdot IAY + \varepsilon \cdot EA - \mu_A \cdot IA \quad \text{(S16)}$$

$$\frac{dSAE}{dt} = \alpha_{AE} \cdot newSAE - bh_A \cdot \mathrm{End}_A \cdot SA - \mu_{AE} \cdot SAE \quad \text{(S17)}$$

$$\frac{dIAE}{dt} = \alpha_{AE} \cdot newIAE - bh_A \cdot \mathrm{End}_A \cdot IAE - \mu_{AE} \cdot IAE \quad \text{(S18)}$$

$$\frac{dnewSAE}{dt} = Eg_{AE} \cdot EgN_{AE} \cdot (SA + ((1 - q) \cdot IA)) - \alpha_{AE} \cdot newSAE - \mu_{AE} \cdot newSAE \quad \text{(S19)}$$

$$\frac{dnewIAE}{dt} = Eg_{AE} \cdot EgN_{AE} \cdot q \cdot IA - \alpha_{AE} \cdot newIAE - \mu_{AE} \cdot newIAE \quad \text{(S20)}$$

The equations for the *Culex* free-living stages are given, followed by the equations for the egg stages. These were compartmentalized as susceptible larvae and pupae, SCLP, young adults (prior to first feeding), SCY, adults, SC, and eggs, SCE, exposed adults, EC and infected adults, IC.

$$\frac{dSCLP}{dt} = bh_C \cdot \text{Hatch}_C \cdot SCE \cdot \left(1 - \frac{SCLP}{NCLP_{max}}\right) - \text{dev}_{CLP} \cdot SCLP - \mu_{CLP} \cdot SCLP \quad \text{(S21)}$$

$$\frac{dSCY}{dt} = \text{dev}_{CLP} \cdot SCLP - \text{wait}_C \cdot SCY - \mu_C \cdot SCY \quad \text{(S22)}$$

$$\frac{dSC}{dt} = \text{wait}_C \cdot SCY - \theta_{CSL} \cdot SC \cdot (IL + IS) - \mu_C \cdot SC \quad \text{(S23)}$$

$$\frac{dEC}{dt} = \theta_{CSL} \cdot SC \cdot (IL + IS) - \epsilon \cdot EC - \mu_C \cdot EC \quad \text{(S24)}$$

$$\frac{dIC}{dt} = \epsilon \cdot EC - \mu_C \cdot IC \quad \text{(S25)}$$

$$\frac{dSCE}{dt} = Eg_{CE} \cdot EgN_{CE} \cdot SC + \delta \cdot Eg_{CE} \cdot EgN_{CE} \cdot IC - bh_C \cdot \text{Hatch}_C \cdot SCE - \mu_{CE} \cdot SCE \quad \text{(S26)}$$

The frequency dependent transmission rates were calculated as follows:

$$\beta_{SLA} = \frac{\text{bite}_A}{NS+NL} \cdot \theta_{SLA} \quad \text{(S27)}$$

$$\beta_{ASL} = \frac{\text{bite}_A}{NS+NL} \cdot \theta_{ASL} \quad \text{(S28)}$$

$$\beta_{SLC} = \frac{\text{bite}_C}{NS+NL} \cdot \theta_{SLC} \quad \text{(S29)}$$

$$\beta_{CSL} = \frac{\text{bite}_C}{NS+NL} \cdot \theta_{CSL} \quad \text{(S30)}$$

The initial conditions for each population were: SS = 360, RS = 240, SL = 140, AL = 60, SAE = 17,946,000, IAE =54,000, SCE = 1000 and all other initial populations were zero.

**Climate-Driven Hatching and Development Rates**

The hatching of both *Aedes* and *Culex* were triggered by rainfall once the temperature exceeded a threshold of 20$^o$C. To simulate the univoltine life cycle (2), seasonal hatching of *Aedes* was triggered early in the rainy season (3-5), once 38 mm cumulative rainfall had fallen during seven days, while the mean daily temperature was greater than 20$^o$C. *Aedes* eggs hatched for nine days (Table S2). These parameters were estimated based on preliminary simulations to represent sufficient rainfall to allow the larvae to hatch following flooding in the *Aedes*, and to ensure that some amount of standing water remained in containers or natural water sources for *Culex* to continue hatching.

The delay (H) in the hatching of *Culex* eggs after the *Aedes* start hatching was set at 7 days (6, 7), provided the temperature is greater than 20$^o$C and there has been greater than 2 mm of cumulative rainfall over three days (to ensure there is standing water present). We simulated the process of *Culex* reproduction throughout the entire year provided the temperature and rainfall thresholds continued to be met.

To simulate the temporal changes in standing water area on which *Culex* can lay their eggs, we used the proportion, $\text{Hatch}_C$, where

$$\text{Hatch}_C = \frac{\text{10 day cumulative rainfall}}{\text{the maximum 10 day cumulative rainfall for that particular year}}. \quad \text{(S31)}$$

$\text{Hatch}_C$ was multiplied by $bh_C$ (the reciprocal of the *Culex* hatching time) to support hatching during the periods

The daily development rates of both *Aedes* and *Culex* larvae and pupae ($\text{dev}_{LP}$ in days$^{-1}$) were modelled as temperature-dependent and were calculated based on the equation (Eq. S2), where $K$ is the daily mean temperature in Kelvin, R is the universal gas constant (1.987 cal·K$^{-1}$·mol$^{-1}$), $K_{C25}$ is the 298.15 Kelvins (25°C), and the species dependent parameters $\rho_{025}$, and HA, HH and TH are given in Rueda *et al*. (8) (Table S3):

$$\text{dev}_{LP} = \frac{\rho_{025} \cdot \frac{K}{K_{C25}} \cdot \exp\left(\frac{HA}{R} \cdot \left(\frac{1}{K_{C25}} - \frac{1}{K}\right)\right)}{1 + \exp\left(\frac{HH}{R} \cdot \left(\frac{1}{TH} - \frac{1}{K}\right)\right)} \quad \text{(S32)}$$

The rate formula developed by Rueda *et al*. (8) was based on data from *Aedes aegypti* and *Culex quinquefasciatus*. To adjust for a difference in species development time, the development rate ($\text{dev}_{ALP}$) for *Aedes mcintoshi* was increased by a factor, $f_{Am}$ (Table S3), as that species develops at a faster rate (6). Similarly, the development rate ($\text{dev}_{CLP}$) for *Culex pipiens* is slower than that for *C. quinquefasciatus* (9) and therefore it was multiplied by $f_{Cp}$ (Table S3) .

The larval/pupal mortality rates for *Aedes* and *Culex* ($\mu_{\text{LP}}$) were derived from the following equation (parameters are defined in Table S4):

$$\phi = \frac{\text{dev}_{\text{LP}}}{\text{dev}_{\text{LP}} + \mu_{\text{LP}}} \quad \text{(S33)}$$

using the genus-specific development rate and survival fractions ($\phi$) obtained from the literature (Table S4). The demographic dynamics of the simulated vector populations (Figure S12) were consistent with patterns of population growth expected.

**Assumptions Made about Host and Vector Populations**

The following assumptions were made about the sheep population:

- Only lambs were imported into the flock.
- Rams were sold upon reaching adulthood such that all adult sheep in the flock were female.
- There was no specific lambing season as farmers in South Africa often manage their sheep to lamb three times every two years (Cordel, pers. comm.).
- Lambs born to recovered ewes had maternal immunity that waned before they mature into sheep.
- No exposed compartment was included among the sheep populations as the incubation period of RVFV in lambs is only 12-20 hours (10).
- If the vaccination scenario is run, only lambs are vaccinated (unless burst is selected), and they mature into vaccinated adult sheep. Further, the vaccine is assumed to be 100% efficacious with no waning of immunity.
- If the vaccine burst scenario is run, 99% of both lambs and adult sheep are vaccinated during the first seven days of July that year.

The following assumptions were made about the *Aedes* and *Culex* mosquito populations:

- The carrying capacity acts via competition and predation on the number of larval and pupae stages.
- Hatching and development were driven by rainfall and temperature as described above (8).
- Larvae and pupae stages were modelled together as one stage.
- Newly metamorphized adult mosquitoes did not take their first blood meal until after the first three days.
- *Aedes* eggs do not hatch in the same hatching period as they were laid as the eggs require a period for embryogenesis and enter a period of dormancy or diapause until the next flood (11).

**Parameters Used in the Model**

The values used for the model parameters were primarily obtained from the literature (Table 4). For those that were not available in the literature we used expert opinion. For six parameters, transovarial transmission, the mortality rate of *Aedes* eggs, *Aedes* and *Culex* bite rates, and *Aedes* and *Culex* carrying capacity, no reliable estimates were available. To identify realistic estimates, we used Latin hypercube sampling to evaluate 80,000 combinations of these six parameters. The results were evaluated by six criteria: mean annual seroprevalence was between 5-40%, the virus persisted for the entire simulation, at least one outbreak “spike” was detected, the ratio of infected *Aedes* eggs remain relatively constant during the simulation and the mean seroprevalence remained relatively constant during the simulation. An outbreak was considered a “spike” if the ratio of infected animals in that year compared to the average number of infected animals in the three years before and after was 2:1 or greater. The population of *Aedes* eggs and mean seroprevalence was determined to be “relatively constant” if the ratio of the mean value from the first half of the simulation to that of the second half of the simulation was within 0.9-1. For the infected *Aedes* eggs we averaged the annual maximum number of eggs. For the seroprevalence we averaged the annual mean seroprevalence. There were three general set of patterns that arose among the six simulations that satisfied the six criteria and these are shown in Figure S13. We evaluated each of them visually and selected the simulation where the outbreak pattern that best matched the historical record of outbreaks in this part of South Africa. The parameter estimates from the exemplar simulation were used to model all simulations presented, except where specific parameters were changed to evaluate the different scenarios.

### Sensitivity Analysis

Three types of sensitivity analyses were conducted. First, we varied one or two parameters at a time and evaluated persistence and mean annual seroprevalence across the 34-year simulation. The parameters included in this first analysis were the transovarial transmission, *Aedes* bite rate, host-to-*Aedes* and *Aedes*-to-host transmission parameters, external incubation rate (EIR) and the proportion of the flock vaccinated.

Second, we used Latin hypercube sampling to explore the model parameter space (12, 13). Fourteen parameters were included in the sensitivity analysis: $\sigma$, $\rho_S$, $\rho_L$, $\mu_A$, $\epsilon$, $\theta_{ASL}$, $\theta_{SLA}$, $\theta_{CSL}$, $\theta_{SLC}$, $\mathrm{bite}_A$, $\mathrm{bite}_C$, $\mu_{AE}$ and $q$, $\mu_C$. The parameters were varied across a range that was 50% above and below the value used in the simulation (or 1.0 if 50% above a proportion was greater than 1.0; see Table 5). The sensitivity analysis consisted of 4000 iterations. Each parameter was evaluated based on its effect on six outcomes: RVFV persistence (years), mean annual seroprevalence, proportion of infected eggs at end of simulation, mean outbreak length, mean outbreak size, and the maximum single outbreak size. These outcomes were calculated for the full 34-year simulation interval. The results were then analyzed using a partial coefficient correlation analysis (12). The third analysis was to explore the effect various parameters had on $R_0$, as described below.

### Estimation of the Basic Reproduction Number, $\boldsymbol{R_0}$

$R_0$ was calculated using the next generation matrix (NGM) approach (14). The first step was to use a version of the model equations with constant rather than varying birth rates and evaluate $R_0$ at a fixed population size. This process was necessary as the carrying capacity parameter in the *Aedes* and *Culex* equations allowed for rapid population changes. This allowed us to solve for $R_0$ at the simulated, constant susceptible populations into which RVFV was introduced. The birth rate necessary to maintain a constant population size are provided in equations S34-S36:

$$\text{birth}_{NSL} = \frac{(NS + NL) \cdot (\mu_L + \text{vax} + g) \cdot (\mu_S + g) \cdot (\mu_S + \text{sold}_S)}{NS \cdot (\mu_S + \text{sold}_S + g) \cdot (\mu_S + \text{vax} + g)} \quad \text{(S34)}$$

$$\text{birth}_A = \frac{\mu_A \cdot (\mu_A + \text{wait}_A) \cdot (\mu_{AE} + \text{bh}_A) \cdot (\mu_{AE} + \alpha_{AE}) \cdot (\mu_{ALP} + \text{dev}_{ALP})}{\text{bh}_A \cdot \text{dev}_{ALP} \cdot \text{wait}_A \cdot \alpha_{AE}} \quad \text{(S35)}$$

$$\text{birth}_C = \frac{\mu_C \cdot (\mu_C + \text{wait}_C) \cdot (\mu_{CE} + \text{bh}_C) \cdot (\mu_{CLP} + \text{dev}_{CLP})}{\text{bh}_C \cdot \text{dev}_{CLP} \cdot \text{wait}_C} \quad \text{(S36)}$$

Briefly, to solve for $R_0$, we first calculated the disease-free equilibrium (DFE) proportions of the host and vector population stages. The vaccinated lamb and sheep populations were set to 0 for the $R_0$ calculation. We then linearized the infectious subsystem using the birth rates calculated as described above. We calculated the Jacobian matrix and decomposed it into the $T$ matrix (the transmission parameters that describes the production of new infections) and $\Sigma$ matrix (the development part that describes changes in state) (14). The NGM with large domain $K_L = -T\Sigma^{-1}$ was calculated and we used the NGM with small domain $K_S$ (using only the rows and columns of $K_L$ belonging to new states-at-infection) to calculate $R_0$. We then solved $R_0$ numerically using set population sizes, the mean population size of sheep and lambs and for each species of vector the 34-year mean population and the mean annual peak population size, calculated across timesteps where there was at least one adult mosquito present. Though the mean peak populations of *Aedes* and *Culex* occurred at different times in the simulation, the estimate of $\boldsymbol{R_0}$ did not allow the incorporation of temporal variation in the populations and therefore we included the mean peak populations from each mosquito species.

We investigated the sensitivity of $R_0$ at fixed host and vector population sizes by varying a single parameter in increments of 25% from 50% below and above the parameter value used in the model (Table S4) while holding the remaining parameters constant. All parameters were included in the sensitivity analysis. We used the 14 parameters that were varied for the Latin hypercube sensitivity analysis (Table S5) to examine the range of $R_0$ values that could be generated as each parameter was varied. For seven of these variables, *Aedes* and *Culex* mortality, bite rates and *Aedes*- and *Culex*-to-host and transovarial transmission fraction, we conducted a sensitivity analysis (parameters were varied from 0.25 and 0.5 times below to above the value used in the simulation) to examine their effects on $R_0$ in populations of *Aedes* and *Culex*, only *Aedes*, or only *Culex* at the mean and mean peak mosquito population levels. To explore the parameter space around transovarial transmission we produced a contour plot of transovarial transmission, *Aedes* bite rate (0.01-0.45) and seroprevalence. Specifically, we investigated the effect of varying TOT with *Aedes* bite rate on mean seroprevalence, long-term (34-year) persistence, short-term ($R_0$ ≥1) persistence, the seasonal $R_0$, and when the seasonal $R_0$ was at unity. The "seasonal" $R_0$ was defined as the value of $R_0$ when transovarial transmission was set at zero to examine whether transovarial transmission was necessary for long-term persistence.

The vaccination threshold that drove RVFV to extinction was identified using simulation.
To explore the extinction timeframe indicated by the vaccination simulations we ran an additional simulation to evaluate the system at 13.5% and 54% vaccination levels. The persistence of RVFV in the host and *Aedes* egg populations was assessed for varying vaccination levels. For the host population, a conservative estimate of greater or equal to one infected host per year indicated persistence. In the *Aedes* egg population, we ran several different simulations varying the initial population of the infected *Aedes* eggs to determine the population threshold required to support viral expansion in the host population. We found that a population of 14,400 infected eggs was sufficient support RVFV expansion. This threshold was used to determine whether there was a sufficient infected *Aedes* egg population to infect hosts in the presence of a sufficiently susceptible host population.

Finally, in addition to continuous vaccination level, we examined the use of burst vaccination, defined as vaccinating 99% of adult sheep and lambs during a seven-day period, once a year (specifically July 1-7, which is in the winter season). Based on the years during which burst vaccination is desired, a vector was created with a daily value of 0 (no burst vaccination) or 1 (burst vaccination) and was fed into the derivative (see below for specific program information).

**Data and Analysis**

We used daily rainfall data from January 1, 1983 to April 30, 2017 from a representative farm in the western Free State (latitude: -28.40, longitude: 26.26; rounded for privacy) extracted from National Oceanic and Atmospheric Administration - Climate Prediction Center (NOAA-CPC) African Rainfall Climatology (ARC) data records (15, 16). The ARC data were produced using a combination of METEOSAT cold cloud duration satellite data and rainfall gauge measurements to produce gridded rainfall estimates at a 0.1° x 0.1° spatial resolution since 1983 (see Figure S1A). The temperature data were collected by the nearest local weather stations maintained under the South African Weather Service (SAWS) at the Glenn College site in Bloemfontein, Free State (latitude: -28.95, longitude: 26.33). The stations (Glen College AWS and Glen College AGR) provided the maximum and minimum temperatures in degrees Celsius. As it was a more complete data set, the Glen College AGR station data were used and the Glen College AWS data were only used if data were missing from the AGR station (SAWS unpublished data, 2017). Temperature data were available from April 1, 1915, to August 2017; however, the dates used were restricted to match the timeframe of the rainfall data (see Figure S1B).

Data analysis was completed in R version 4.1.2 (17) using Rstudio (18). The package zoo 1.8-8 (19) was used to calculate the cumulative rainfall data, the ode solver was used from the deSolve 1.28 package (20), the climate data and other daily data (e.g. the hatching triggers, burst vaccination) were introduced into the simulation through the use of the approxfun() function (deSolve) and the figures were made using ggplot2 3.3.2 (21). The lhs 1.1.1 package (22) was used to produce the Latin hypercube for the sensitivity analysis. The resultant data were analyzed by calculating the partial correlation coefficients using the pcc() function from the sensitivity 1.22.1 package (23). wxMaxima 5.43.0 (24) was used to conduct the $R_0$ analysis.

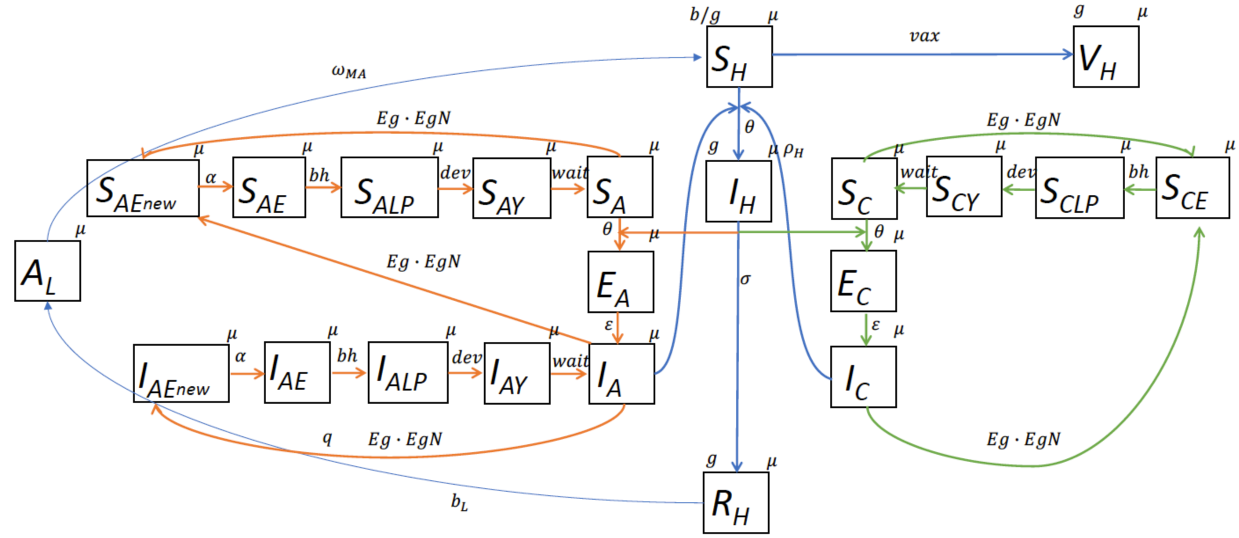


**Fig. S1** A diagram of the mathematical model. The infectious states of Susceptible ($S$), Exposed ($E$), Infected ($I$), Maternal Antibodies ($MA$) and Vaccinated ($V$) are represented for the host ($H$), *Aedes* vectors ($A$) and *Culex* vectors ($C$). All of the life stages of are represented for the two mosquito populations, including susceptible and infected *Aedes* eggs ($S_{AE}$, $I_{AE}$), larvae/pupae ($S_{ALP}$, $I_{ALP}$), young (non-feeding) adults ($S_{AY}$, $I_{AY}$) and adult *Aedes* ($S_A$, $I_A$) as well as susceptible *Culex* eggs ($S_{CE}$), larvae/pupae ($S_{CLP}$), young (non-feeding) adults ($S_{CY}$) and adult *Culex* ($S_C$). *Aedes* eggs are initially laid in a compartment of new eggs that require desiccation ($S_{AEnew}$, $I_{AEnew}$) before moving to the infected egg compartments that can hatch ($S_{AE}$, $I_{AE}$). H represents both developmental stages in sheep, (adult sheep or lambs) while $L$ only represents lambs. Host demographics and losses through sale are not indicated in the diagram.

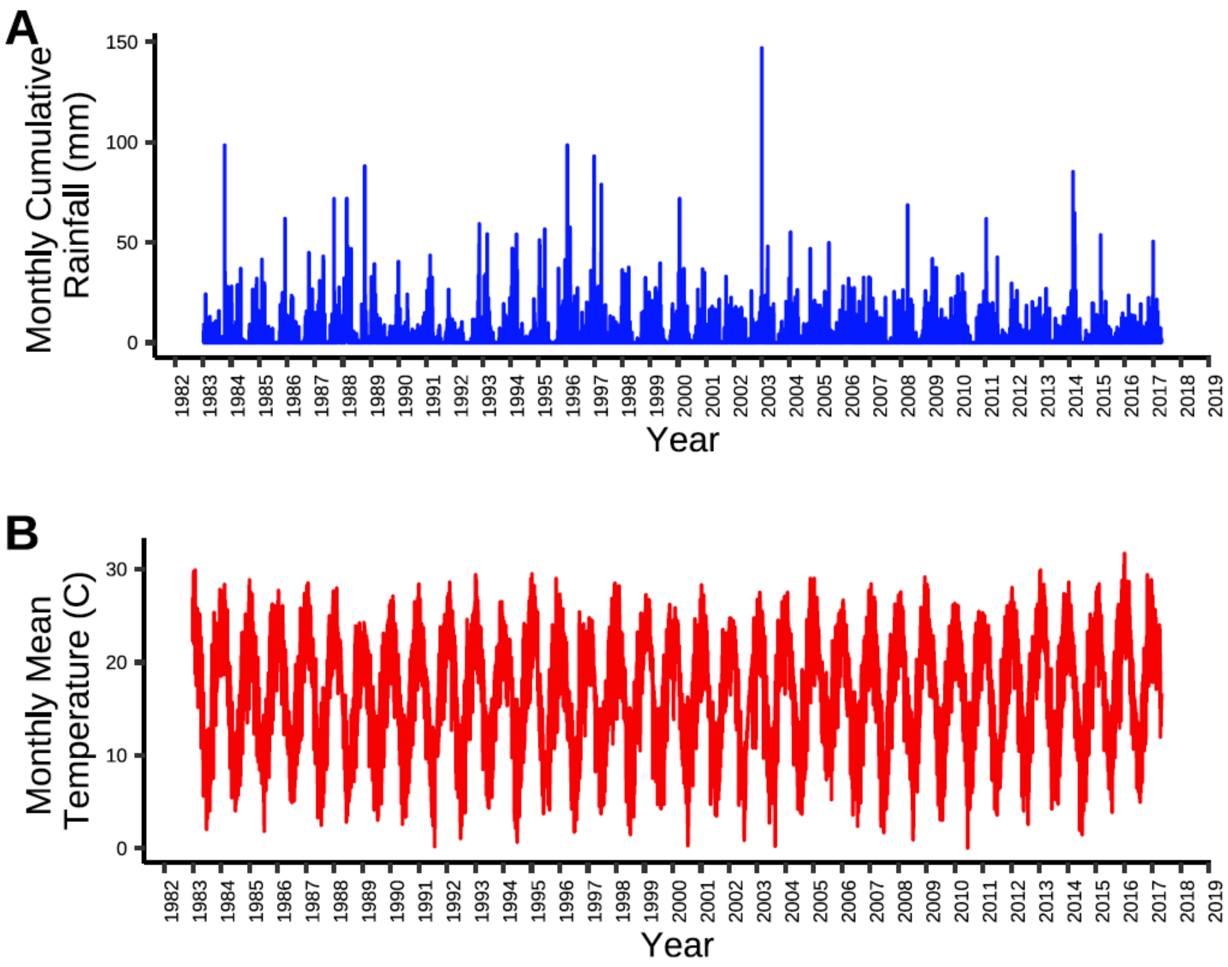


**Fig. S2** Monthly A) total rainfall and B) mean temperature by year from 1983-2017.

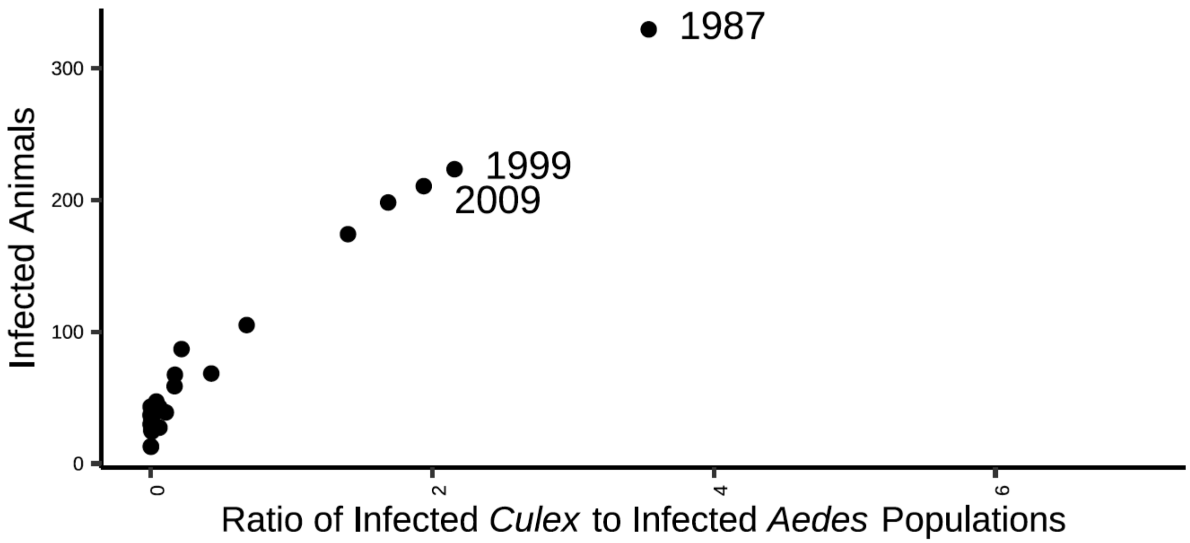


**Fig. S3** The total number of infected hosts by the peak population sizes of infected Culex and Aedes for each year of the simulation. The year of the outbreak is indicated for all outbreaks with a ratio of infected Culex and Aedes >2.

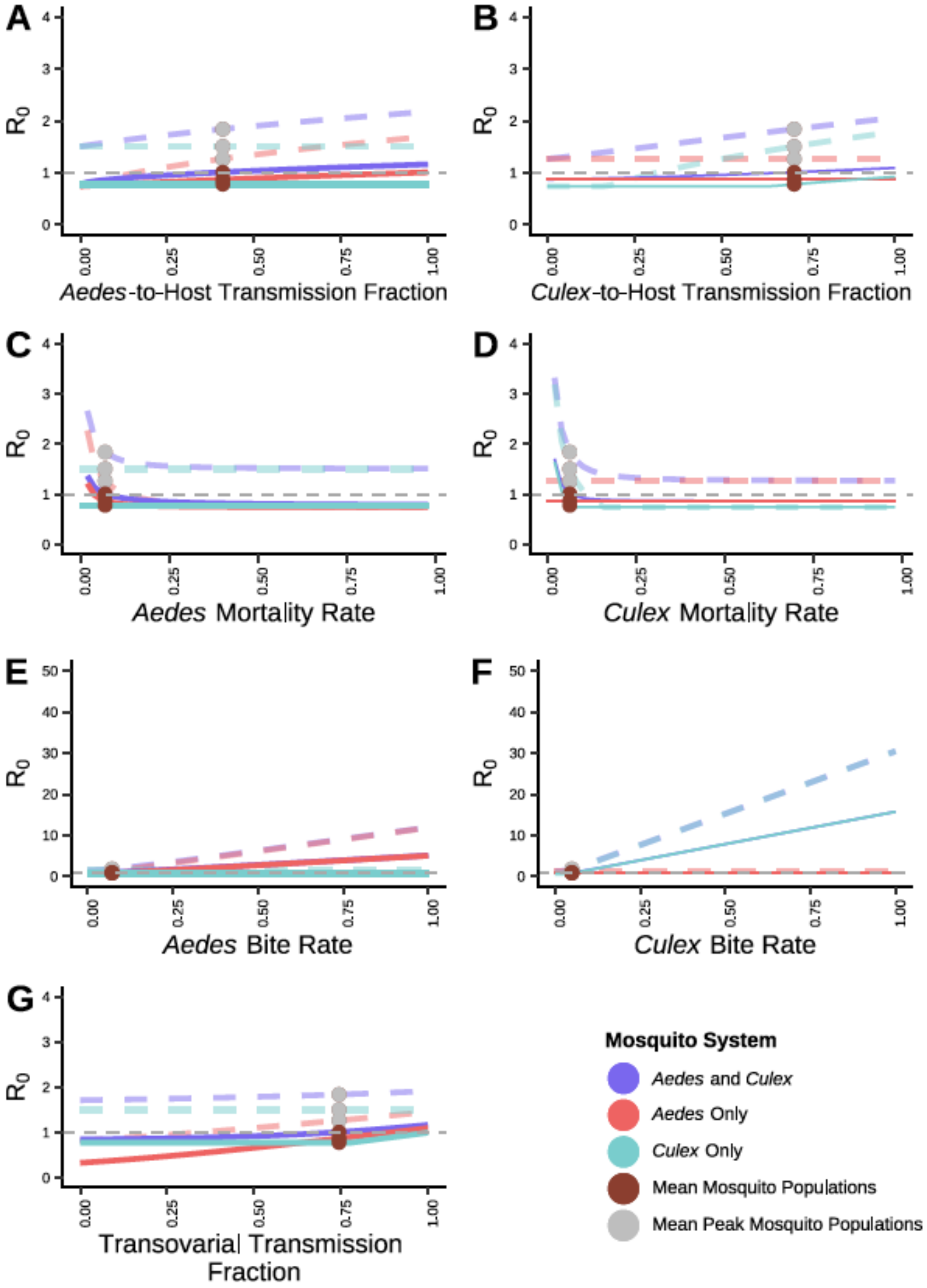


**Fig. S4** Sensitivity analysis of $R_0$ illustrate how $R_0$ changes in relation to A) Aedes-host transmission fraction; B) Culex-host transmission fraction; C) the Aedes mortality rate; D) Culex mortality rate; E) Aedes bite rate; F) Culex bite rate and G) transovarial transmission fraction. The red line represents the system with Aedes mosquitoes only, the teal line represents the system with Culex mosquitoes only and the purple line represents the system with both Aedes and Culex mosquitoes. The solid lines indicate $R_0$ at the mean mosquito population, whereas the dashed lines represent the value of $R_0$ using the mean peak mosquito population. The brown dots indicate the $R_0$ at the parameter value used in our simulation. The dashed gray line indicates $R_0$ equal to one.

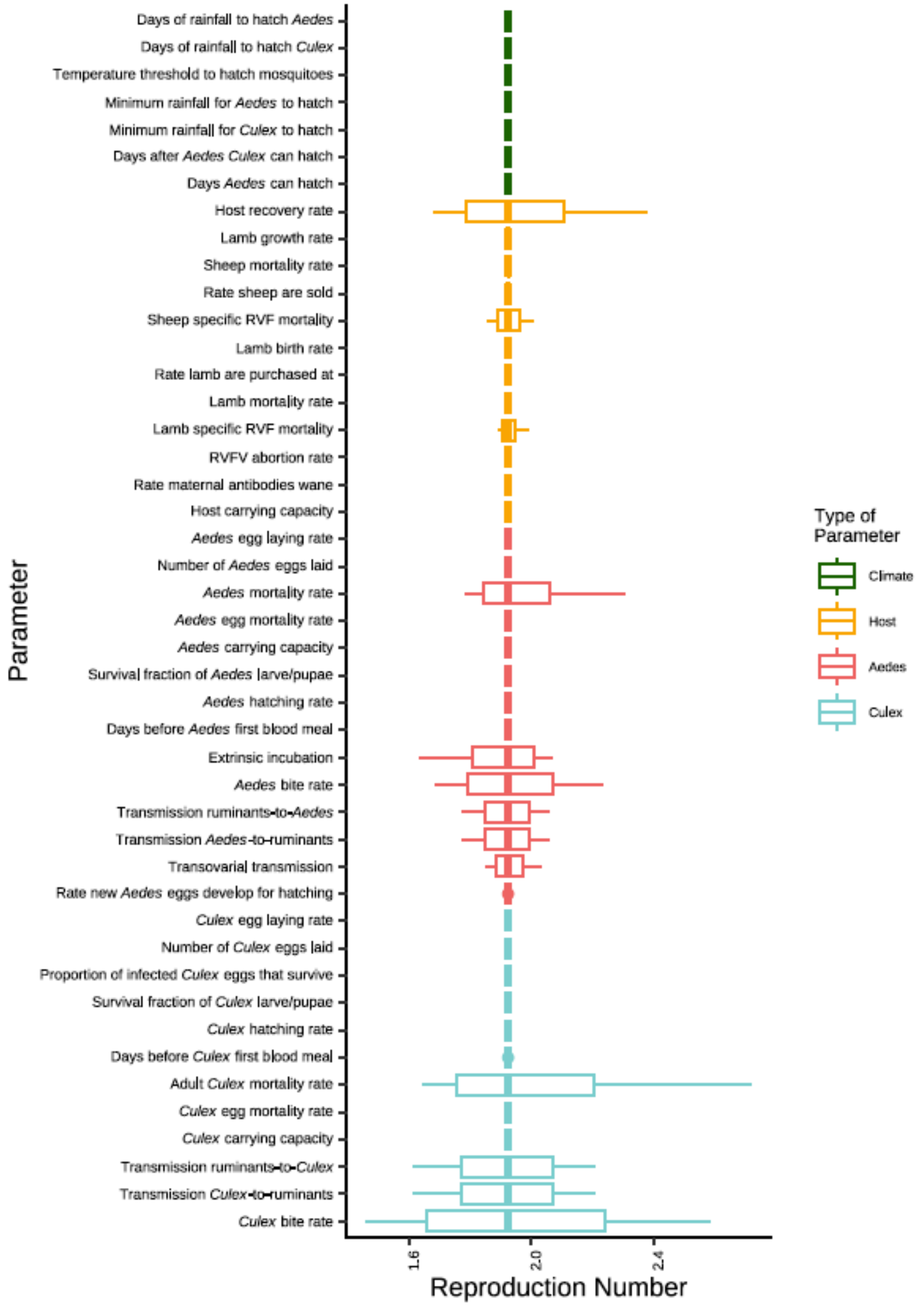


**Fig. S5** The value of $R_0$ across a range of values for each parameter used in the model and calculated at a fixed vector and host population size. The parameters are characterized as those impacting the hatching of the mosquitoes (climate [green]), and those impacting the populations of the hosts (orange), *Aedes* (red) and *Culex* (light blue).

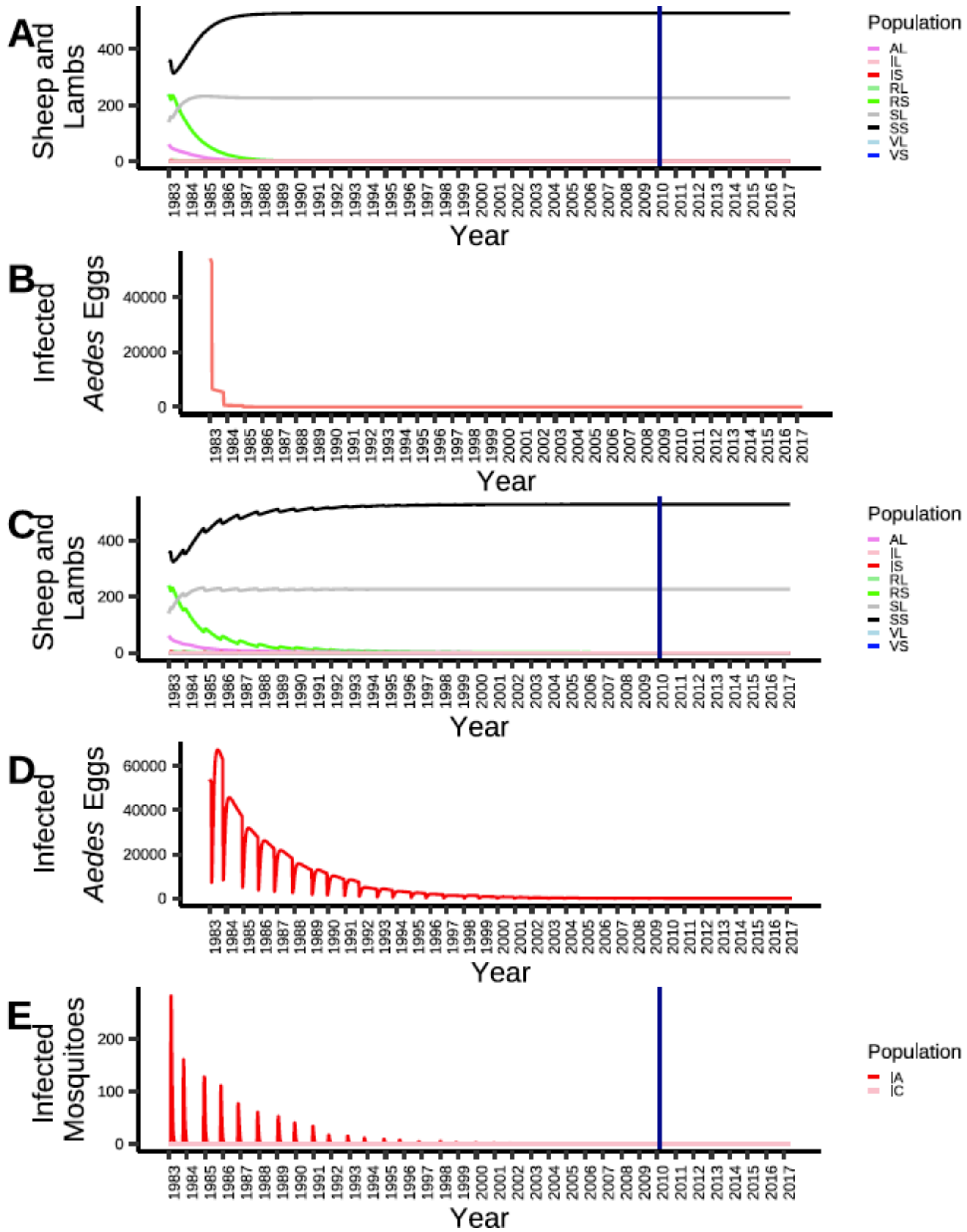


**Fig. S6** The dynamics of RVFV in sheep (A) and infected *Aedes* egg (B) populations in a system with no transovarial transmission. The dynamics of RVFV in sheep (C), infected *Aedes* egg (D) and infected mosquito (E) populations in a system with no horizontal transmission.

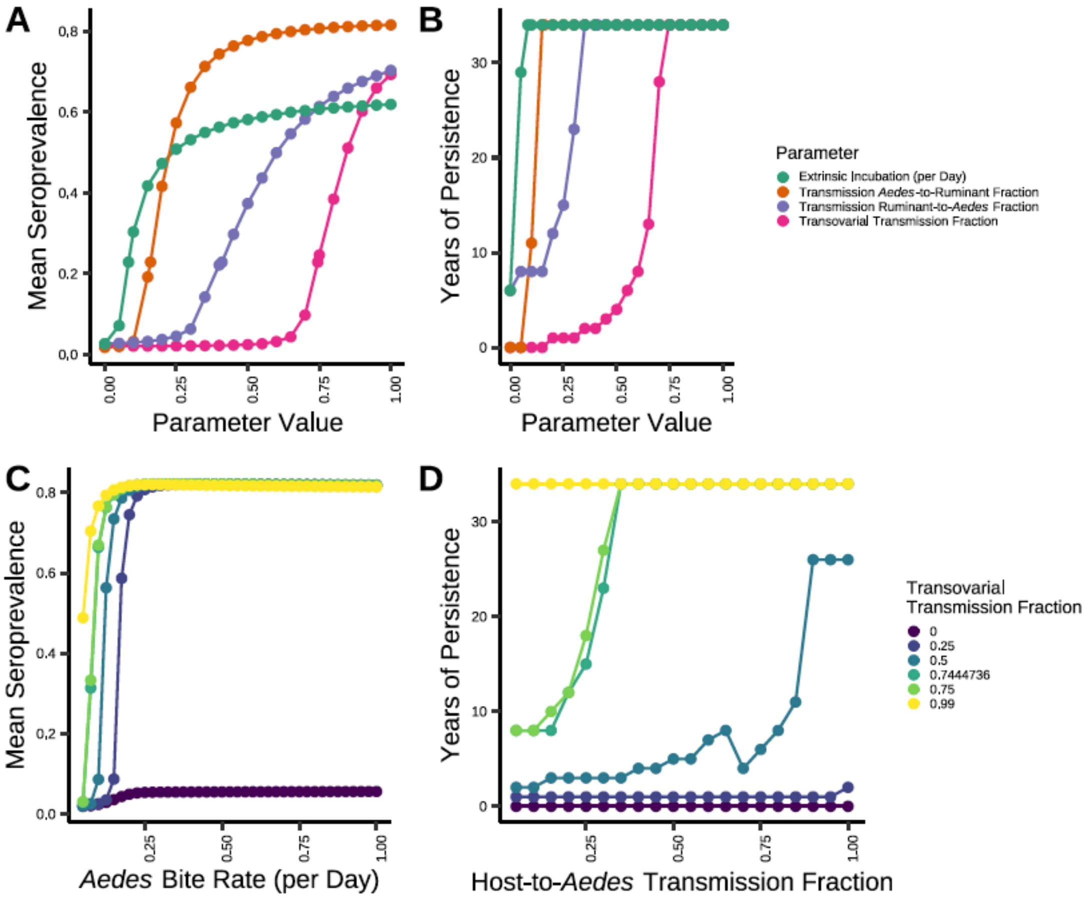


**Fig. S7** The sensitivity of the mean seroprevalence (A) and persistence (B) of the system to the transovarial transmission fraction, horizontal transmission parameters, *Aedes* bite rate and the extrinsic incubation rate. The seroprevalence and persistence of RVFV in the system are more sensitive when considering the parameters together, e.g., sensitivity of seroprevalence to the *Aedes* bite rate and transovarial transmission fraction (C) and persistence to the host-to-vector and transovarial transmission fractions (D).

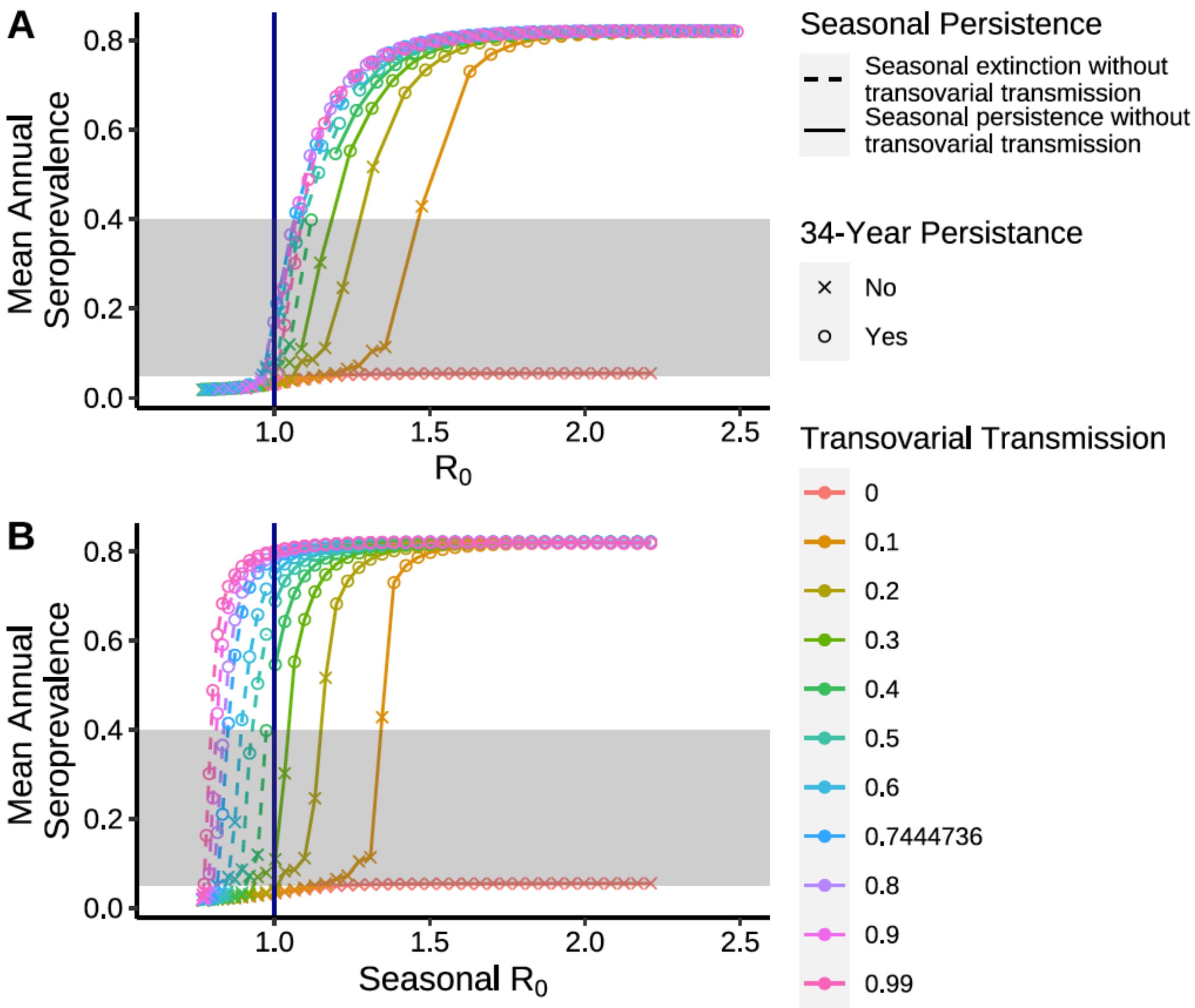


**Fig. S8** Changes in mean annual seroprevalence and $R_0$ as transovarial transmission and *Aedes* bite rate (0.01-0.45 bites per day) changes. Persistence occurs once $R_0$ crosses unity (top) with mean annual seroprevalences in the desired range (5-40%, shaded area) obtained for transovarial transmission fraction values > 0.5. However, for these parameter ranges, the seasonal $R_0$ (i.e., the estimated $R_0$ excluding the contribution of transovarial transmission) is < 1 (bottom). When the seasonal $R_0$ exceeds one, this leads to large outbreaks and a lack of long-term persistence. Thus, achieving long term persistence and observed seroprevalences appears to require limited seasonal spread and an overwintering mechanism via transovarial transmission which boosts $R_0$ above unity. Simulations with long-term RVFV persistence in the host population (the final year of the simulation) are indicated by points, whereas "x" indicates RVFV did not persist. Simulations with seasonal persistence (seasonal $R_0$ is greater than unity) are indicated by the dashed lines and those that did not have seasonal persistence are indicated with a solid line. The blue line indicates $R_0$ at unity.

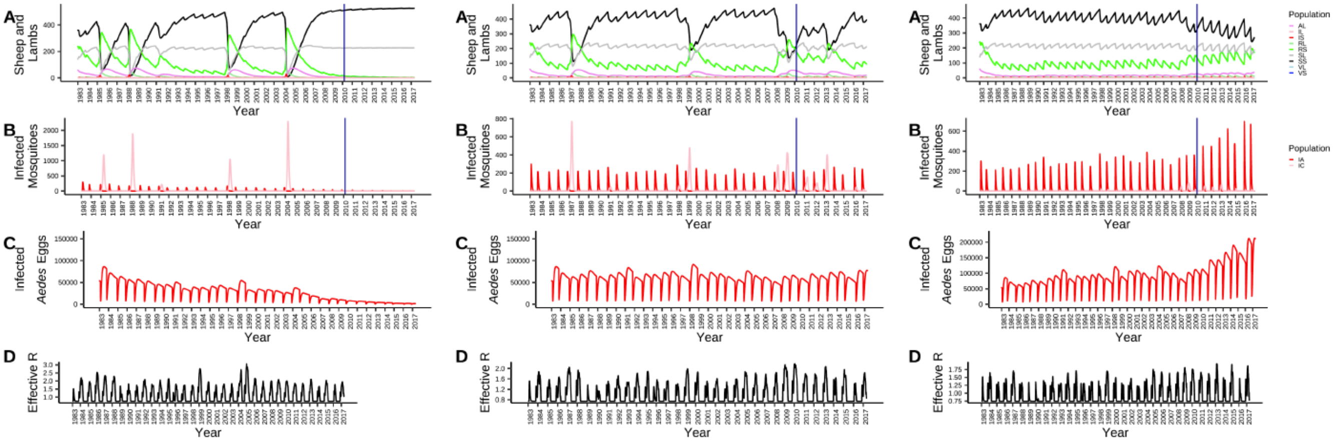


**Fig. S9** The full 34-year simulation of A) hosts, B) infected mosquitoes, C) infected *Aedes* eggs and D) the effective reproduction number using 75% of the *Culex* mortality rate (left), the *Culex* mortality rate at the value used for the model simulation (middle) and 125% of the *Culex* mortality rate (right). AL = lambs with maternal antibodies; IL, IS, IA, IC, and IAE = infected lambs, sheep, *Aedes*, *Culex* and *Aedes* eggs, respectively; RL and RS = recovered lambs and sheep, respectively; SL, SS, SA, and SC = susceptible lambs, sheep, adult *Aedes* and adult *Culex*, respectively. VL and VS = vaccinated lambs and sheep, respectively.

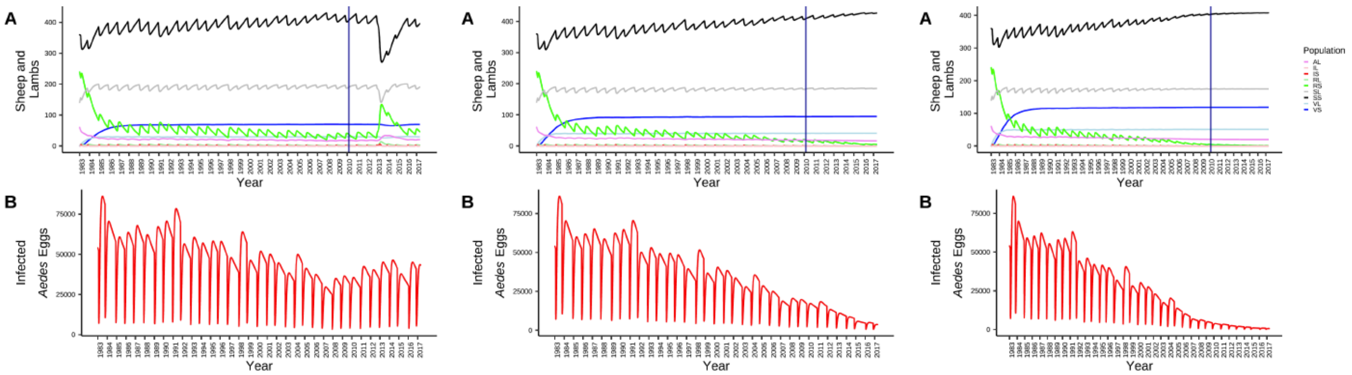


**Fig. S10** A full 34-year simulation of A) hosts and B) infected *Aedes* eggs when Left: the proportion of animals vaccinated is maintained at 13.5%, Center: the proportion of animals vaccinated is maintained at 18% and Right: the proportion of animals vaccinated is maintained at 54%. AL = lambs with maternal antibodies; IL, IS, IA, IC, and IAE = infected lambs, sheep, *Aedes*, *Culex* and *Aedes* eggs, respectively; RL and RS = recovered lambs and sheep, respectively; SL, SS, SA, and SC = susceptible lambs, sheep, adult *Aedes* and adult *Culex*, respectively. VL and VS = vaccinated lambs and sheep, respectively.

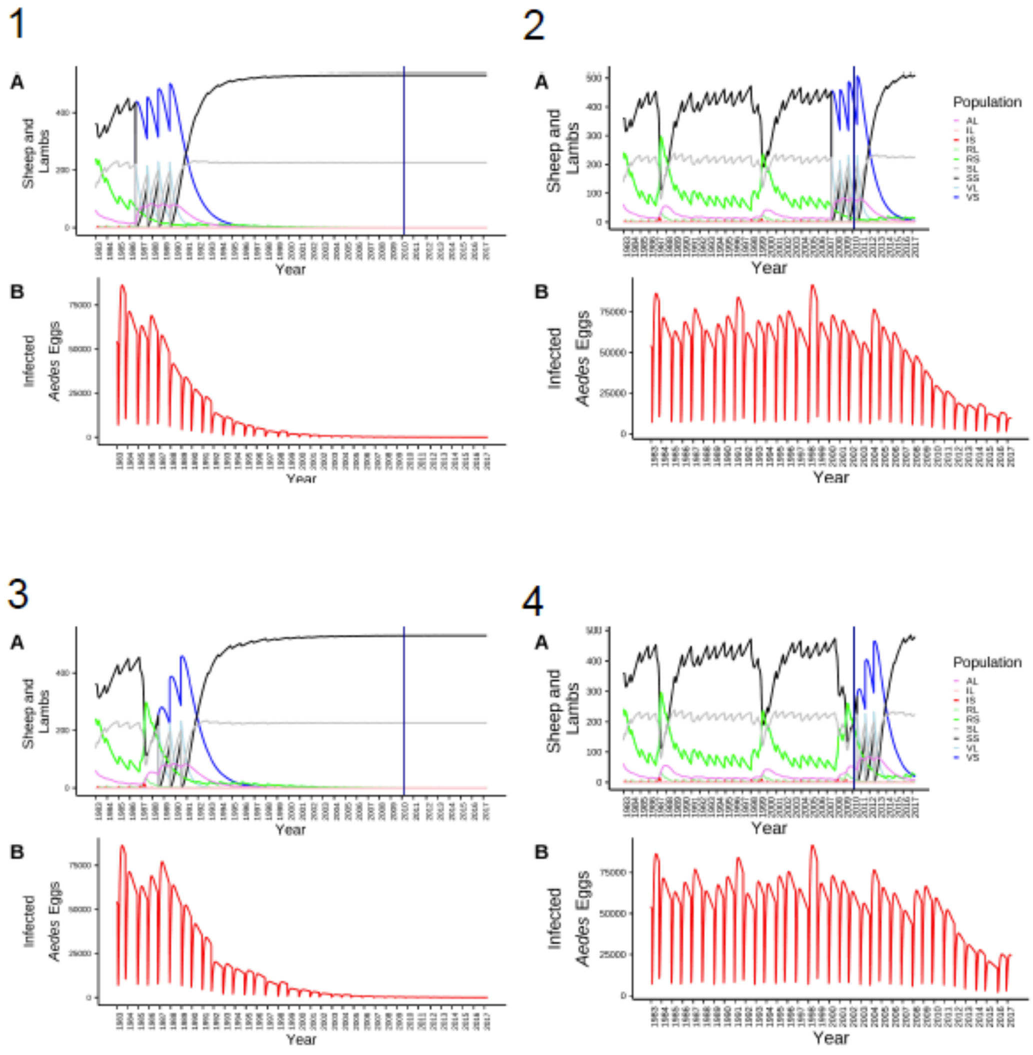


**Fig. S11** A full 34-year simulation of A) hosts and B) infected *Aedes* eggs when “burst” vaccination is used. Burst vaccination is when 99% of hosts are vaccinated on July 1-7 of a given year and there is no vaccination the rest of the year. 1: Right: Burst vaccination is used outside of an outbreak period and early in the simulation and required more years of vaccination (1986, 1987, 1988 and 1989). 2: Burst vaccination is used outside of an outbreak period and late in the simulation and required more years of vaccination (2007, 2008, 2009, 2010). 3: Early in the simulation and immediately after the 1987-88 outbreak burst vaccination is used for three years and to drive RVFV to extinction (1988, 1989, 1990). 4: Burst vaccination combined with a natural outbreak, requires fewer years of vaccination, even later in the simulation (2010, 2011, 2012).

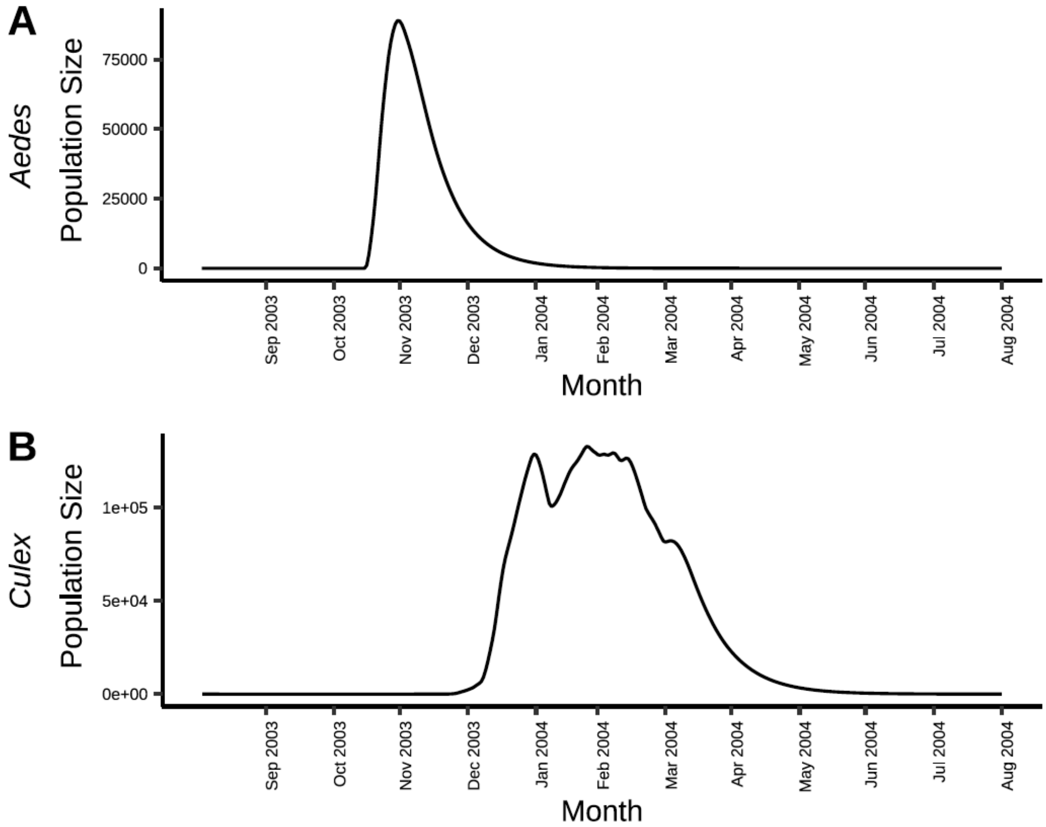


**Fig. S12** The total population of adult A) *Aedes* and B) *Culex* mosquitoes over a randomly selected year (September 2003- August 2004).

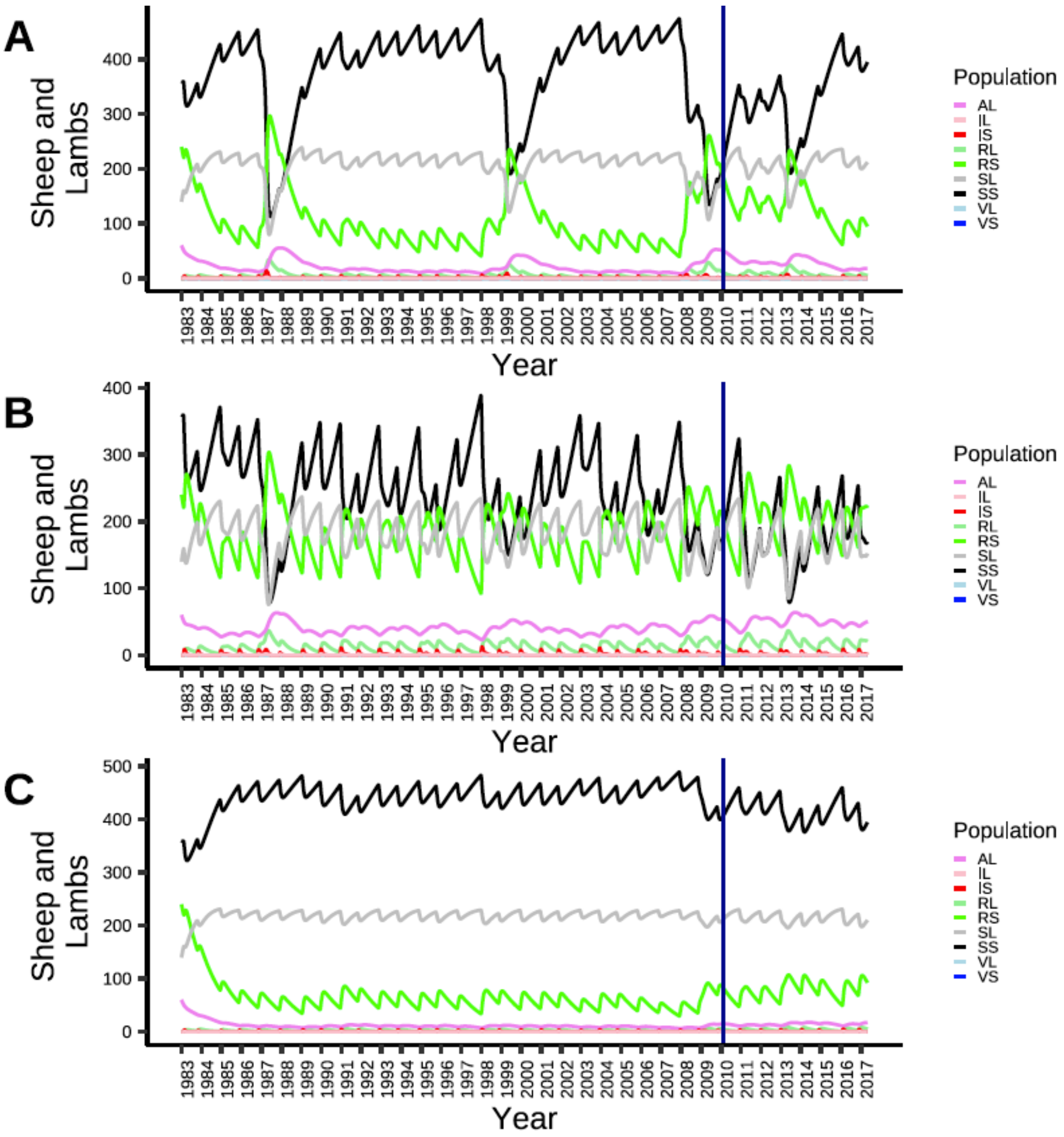


**Fig. S13** The simulations to provide parameter values for the six unknown parameters and satisfied the assessment criteria had three broad patterns A) large outbreaks with intervening periods of low-level transmission, B) high rates of annual transmission resulting in frequent large outbreaks, or C) low-level transmission with occasional and small outbreaks.

**Table S1.** Mean annual seroprevalence, mean annual maximum host-vector ratio, and the mean proportion of infected mosquitoes and eggs during the entire simulation.

| Factor | Mean (Range) |
|---|---|
| Mean annual seroprevalence | 22.9% (10.5-58.6%) |
| Mean annual maximum host-vector ratio | 1:180 (1:51-1:286) |
| Mean annual proportion of infected *Aedes* eggs | 0.003 (0.002-0.003) |
| Mean annual proportion of infected adult *Aedes* | 0.007 (0.002-0.04) |
| Mean annual proportion of infected adult *Culex* | 0.002 (0-0.035) |

**Table S2.** Definition of the state variables used in the model.

| Variable | Population represented |
|---|---|
| Sheep and Lambs | |
| AL | Lambs with temporary maternal immunity |
| SL | Susceptible lambs |
| IL | Infected lambs |
| RL | Recovered lambs |
| VL | Vaccinated lambs |
| SS | Susceptible sheep |
| IS | Infected sheep |
| RS | Recovered sheep |
| VS | Vaccinated sheep |
| *Aedes Mosquitoes* | |
| SA | Susceptible adult *Aedes* |
| EA | Exposed adult *Aedes* |
| IA | Infected adult *Aedes* |
| SAY | Susceptible young *Aedes* that have not had their first blood meal |
| IAY | Infected young *Aedes* that have not had their first blood meal |
| SALP | Susceptible *Aedes* larvae/pupae |
| IALP | Infected *Aedes* larvae/pupae |
| newSAE | Newly laid susceptible *Aedes* eggs that cannot hatch yet |
| SAE | Susceptible *Aedes* eggs that can hatch |
| newIAE | Newly laid infected *Aedes* eggs that cannot hatch yet |
| IAE | Infected *Aedes* eggs that can hatch |
| *Culex* Mosquitoes | |
| SC | Susceptible adult *Culex* |
| EC | Exposed adult *Culex* |
| IC | Infected adult *Culex* |
| SCY | Susceptible young *Culex* that have not had their first blood meal |
| SCLP | Susceptible *Culex* larvae/pupae |
| SCE | Susceptible *Culex* eggs |

**Table S3.** Parameter values for the climate forcing to drive the hatching and development of the mosquitoes as described in the text.

| Description | Value | Unit | Reference |
|---|---|---|---|
| Number of days that *Aedes* eggs can hatch | 9 | days | (27) |
| Number of cumulative days of rainfall needed to initiate hatching of *Aedes* eggs | 7 | days | See main text |
| Minimum amount of rainfall over 7 days, above which *Aedes* can hatch | 38 | mm | See main text |
| Minimum temperature above which both *Aedes* and *Culex* eggs can hatch | 20 | Celsiu s | (28) |
| Number of cumulative days of rainfall needed to initiate and maintain hatching of *Culex* eggs | 3 | days | See main text |
| Minimum amount of rainfall over the 3 days, above which *Culex* can hatch | 2 | mm | See main text |
| Number of days after the *Aedes* eggs start hatching that the *Culex* can start hatching (H) | 7 | days | (27) |

**Table S4.** Parameter values for calculating the larval development rates (Eq. S2). Rueda *et al.* (8) confirmed the Sharpe & DeMichele model (10) with high temperature inhibition with laboratory raised *Aedes* and *Culex* larvae. Their model was built to estimate the development rate per day as the temperature changed (Eq. S2).

| Variable | Value | Description | Reference |
|---|---|---|---|
| $\rho_{025_A}$ | 0.1546 | Effect of temperature on *Aedes* larval development estimated by a linear regression | (8) |
| $HA_A$ | 33255.57 | Effect of temperature on *Aedes* larval development estimated by a linear regression | (8) |
| $TH_A$ | 301.67 | Effect of temperature on *Aedes* larval development estimated by a linear regression | (8) |
| $HH_A$ | 50543.49 | Effect of temperature on *Aedes* larval development estimated by a linear regression | (8) |
| $f_{Am}$ | 1.392704 | Factor by which *Aedes mcintoshi* develops faster than *Culex quinquefasciatus* | Expert opinion |
| $\rho_{025_C}$ | 0.21945 | Effect of temperature on *Culex* larval development estimated by a linear regression | (8) |
| $HA_C$ | 28049.98 | Effect of temperature on *Culex* larval development estimated by a linear regression | (8) |
| $TH_C$ | 298.6 | Effect of temperature on *Culex* larval development estimated by a linear regression | (8) |
| $HH_C$ | 35362.18 | Effect of temperature on *Culex* larval development estimated by a linear regression | (8) |
| $f_{Cp}$ | 0.7650069 | Factor by which *Culex pipiens* develops faster than *Culex quinquefasciatus* | Expert opinion |

**Table S5.** Parameter values for the SEIR model of mosquitoes, sheep, and lambs (Eqs. S4-S33).

| Parameter | Value | Description | Unit | Reference |
|---|---|---|---|---|
| $\sigma$ | 0.25 | Recovery rate | days$^{-1}$ | (12) |
| $g$ | 0.005 | Maturation rate of lambs to sheep | days$^{-1}$ | (13) |
| $\mu_S$ | 0.00014 | Sheep mortality rate | days$^{-1}$ | Rostal *et al* unpublished |
| $sold_S$ | 0.002 | Percent of sheep sold | days$^{-1}$ | Set to maintain sheep population constant |
| $\rho_S$ | 0.063 | RVFV specific mortality rate in sheep | days$^{-1}$ | (14) |
| $b_L$ | 0.004 | Birth rate in sheep | days$^{-1}$ | (15) |
| $buy_L$ | 0.508 | Number of lambs purchased | days$^{-1}$ | Rostal *et al* unpublished |
| $\mu_L$ | 0.00004 | Lamb mortality rate | days$^{-1}$ | Rostal *et al* unpublished |
| $\rho_L$ | 0.751 | RVFV specific mortality rate in lambs | days$^{-1}$ | (16) |
| *Abort* | 0.9 | Abortion fraction among pregnant ewes infected with RVFV | No units | (14) |
| $\omega_{MA}$ | 0.008 | Maternal antibodies Waning rate | days$^{-1}$ | (17) |
| vax.prop | 0 | Proportion of the sheep and lamb populations desired to be vaccinated | No units | Set by user as needed |
| vax | 0 | $\frac{\text{vax.prop} \cdot (mu_L + g)}{(1 - \text{vax.prop})}$ | days$^{-1}$ | Calculated from vax.prop |
| $vax_{scheme.l}$ | [1,0] | Vector to indicate when to apply burst vaccination on adult sheep | No units | Set based on the selection of burst vaccination by user |
| $vax_{scheme.s}$ | [1,0] | Vector to indicate when to apply burst vaccination on adult sheep | No units | Set based on the selection of burst vaccination by user |
| $NL_{max}$ | 800 | Herd size maintained by the carrying capacity acting on lambs | No units | Rostal *et al* unpublished |
| $Eg_{AE}$ | 0.333 | Egg laying Rate for *Aedes circumluteolus* | days$^{-1}$ | (18) |
| $EgN_{AE}$ | 34.583 | Number of female eggs per adult Asian *Aedes lineatopennis* | No units | (19) |
| $\mu_A$ | 0.07 | *Aedes* mortality rate | days$^{-1}$ | (7, 20) |
| $\mu_{AE}$ | 0.0008378592 | *Aedes* egg mortality rate | days$^{-1}$ | See main text above for estimation process |
| $NALP_{max}$ | 223,880.6 | Carrying capacity acting on the number of *Aedes* larvae and pupae that can survive in the available habitat | No units | See main text above for estimation process |
| $\phi_A$ | 0.36 | Fraction of *Aedes mcintoshi* larvae/pupae that survive | No units | (5) |
| $bh_A$ | 0.23 | Rate that *Aedes* eggs hatch based the approximately 5-day interval between | days$^{-1}$ | (6) |

| Parameter | Value | Description | Unit | Reference |
|---|---|---|---|---|
| | | flooding and initial larval detection in the pan | | |
| $\alpha_{AE}$ | 0.033 | The rate at which newly laid eggs desiccate and become able to hatch | days$^{-1}$ | Set to prevent newly laid eggs from hatching during the same annual hatching period in which they were laid |
| $\text{dev}_{ALP}$ | 0.068 | Mean development rate of *Aedes* - calculated daily based on ambient temperature (Eq. S2) | days$^{-1}$ | (8) |
| $\text{wait}_{A}$ | 0.333 | Reciprocal of the number of days (3) it takes before a newly emerged *Aedes* takes its first bite (delay due to mating, host-seeking etc.) | days$^{-1}$ | (21-23) |
| $\mu_{ALP}$ | 0.12 | Mean mortality rate in *Aedes* larvae/pupae on days when there was at least 1 susceptible or infected *Aedes* larvae/pupae - calculated daily based on ambient temperature (Eq. S3) | days$^{-1}$ | Estimated from $\text{dev}_{ALP}$ |
| $\text{bite}_{A}$ | 0.07108191 | *Aedes* bite rate on the flock of sheep | days$^{-1}$ | See main text above for estimation process |
| $\epsilon$ | 0.083 | Extrinsic incubation rate in mosquitoes | days$^{-1}$ | (24) |
| $\theta_{ASL}$ | 0.41 | Probability of transmission to *Aedes lineatopennis/mcintoshi* from the host | No units | (25, 26) |
| $\theta_{SLA}$ | 0.16 | Probability of transmission to the host from *Aedes lineatopennis/mcintoshi* | No units | (25) |
| $q$ | 0.7444736 | Transovarial transmission fraction | No units | See main text above for estimation process |
| $\text{End}_{A}$ | [1,0] | When the value is one, *Aedes* eggs hatch; represents the univoltine hatching of *Aedes* | No units | Estimated by climate thresholds |
| $Eg_{CE}$ | 0.333 | Egg laying rate of *Culex* | days-1 | (24) |
| $EgN_{CE}$ | 118 | Mean number of female eggs laid by *Culex* | No units | (9) |
| $\delta$ | 0.8 | Fraction of RVFV infected *Culex* that lay eggs | No units | (27) |
| $\phi_{C}$ | 0.291 | Fraction of *Culex* that survive larval/pupal stages | No units | (9) |
| $bh_{C}$ | 0.667 | Reciprocal of the *Culex* hatching time | days$^{-1}$ | (9) |
| $\text{dev}_{CLP}$ | 0.06 | Mean development time for *Culex* pupae/larvae - calculated daily based on ambient temperature (Eq. S2) | days$^{-1}$ | (8) |
| $\text{wait}_{C}$ | 0.333 | Reciprocal of the number of days (3) it takes before a newly emerged *Culex* | days$^{-1}$ | (21-23) |

| Parameter | Value | Description | Unit | Reference |
|---|---|---|---|---|
| | | takes its first bite (delay due to mating, host-seeking etc.) | | |
| $\mu_C$ | 0.063 | Adult *Culex* mortality rate | $days^{-1}$ | (7, 24, 28) |
| $\mu_{CLP}$ | 0.145 | Mean mortality rate in *Culex* larvae/pupae - calculated daily based on ambient temperature (Eq. S3) | $days^{-1}$ | Estimated from $\mathrm{dev}_{CLP}$ |
| $\mu_{CE}$ | 0.146 | *Culex* egg mortality rate - calculated using $\mu_{CE} = bh_C \cdot \frac{1-\psi_C}{\psi_C}$ | $days^{-1}$ | Estimated as per the equation |
| $NCLP_{max}$ | 110,013.6 | Carrying capacity acting on the number of *Culex* larvae and pupae that can survive in the available habitat | No units | See main text above for estimation process |
| $\theta_{CSL}$ | 0.71 | Probability of transmission to *Culex* from the host | No units | (25, 26, 29-32) |
| $\theta_{SLC}$ | 0.31 | Probability of transmission to the host from *Culex* | No units | (25, 26, 32) |
| $\mathrm{bite}_C$ | 0.04952358 | *Culex* bite rate on the sheep | $days^{-1}$ | See main text above for estimation process |
| $\mathrm{Hatch}_C$ | 0-1 | When the value is greater than zero, *Culex* eggs hatch at the specified proportion | No units | Estimated by climate thresholds |

**Table S6** Ranges of parameters used in the Latin hypercube and $\boldsymbol{R_0}$ sensitivity analyses.

| Parameter | Range used in Latin hypercube |
|---|---|
| $\sigma$ | 0.125-0.375 |
| $\rho_S$ | 0.0315-0.0945 |
| $\rho_L$ | 0.3755-1.0 |
| $\epsilon$ | 0.0415-.01245 |
| $\mathrm{bite}_A$ | 0.06-0.18 |
| $\theta_{ASL}$ | 0.205-0.615 |
| $\theta_{SLA}$ | 0.08-0.24 |
| $\mu_A$ | 0.035-0.105 |
| $\mu_{AE}$ | 0.00005-.0015 |
| $q$ | 0.35-1.0 |
| bite$_C$ | 0.0175-0.0525 |
| $\theta_{CSL}$ | 0.35-1.0 |
| $\theta_{SLC}$ | 0.0175-0.155 |
| $\mu_C$ | 0.0315-.0945 |
